# Interactions between two QTLs for time to anthesis on spike development and fertility in wheat

**DOI:** 10.1101/2020.11.11.378208

**Authors:** Priyanka A. Basavaraddi, Roxana Savin, Luzie U Wingen, Stefano Bencivenga, Alexandra M. Przewieslik-Allen, Simon Griffiths, Gustavo A. Slafer

## Abstract

Earliness per se (*Eps*) genes are reported to be important in fine-tuning flowering time in wheat independently of photoperiod (*Ppd*) and vernalisation (*Vr*n). Unlike *Ppd* and *Vrn* genes, *Eps* have relatively small effects and their physiological effect along with chromosomal position are not well defined. We evaluated eight lines derived from crossing Paragon and Baj (late and early flowering respectively), vernalisation insensitive, to study the detailed effects of two newly identified QTLs, *Eps-7D* and *Eps-2B* and their interactions under field conditions. The effect of both QTLs were minor but their effect was modulated by the allelic status of the other. While the magnitude of effect of these QTLs on anthesis was similar, they are associated with very different profiles of pre-anthesis development which also depends on their interaction. *Eps-7D* affected both duration before and after terminal spikelet while not affecting final leaf number (FLN) so *Eps-7D*-*early* had a faster rate of leaf appearance. *Eps-2B* acted more specifically in the early reproductive phase and slightly altered FLN without affecting the leaf appearance rate. Both *Eps-7D* and *2B* affected the spike fertility by altering the rate of floret development and mortality. The effect of the latter was very small but consistent in that the -*late* allele tended to produced more fertile florets.

## Introduction

Wheat adaptation has been possible by coarse tuning time to anthesis^1,2^. Moreover, fine tuning time to anthesis is relevant to improve adaptation and performance within a specific environment^3–5^. The possibility of coarse and fine tuning of time to anthesis in wheat (and other winter cereals) has helped to maximise yield because of optimised anthesis time^6,7^; and will be instrumental in adapting better to climate change^8,9^.

Wheat development from sowing to anthesis encompasses various stages that make up two major component phases: time from sowing to terminal spikelet (TS), combining the vegetative and early reproductive phases of leaf and spikelet initiation, and time from TS to anthesis, i.e. the late reproductive phase (LRP) of floret initiation and survival^10^. Duration of each of these phases plays a key role in determining the generation and degeneration of various organs (shoots, spikes, spikelets, florets and grains) that define components of yield^11,12^. It has been hypothesised that developmental phases prior to anthesis can be adjusted with little or no changes in time to anthesis to keep improving adaptation along with stable or improved yield^13,14^

Wheat responds to various environmental stimuli like vernalisation and photoperiod and genetic factors responsible for these sensitivities are mainly *Vrn* and *Ppd* genes, respectively^1^. Responses to photoperiod and vernalisation sensitivities can be found in all the three (vegetative, early and later reproductive) phases of wheat development^10,13,15,16^. The effects of different *Ppd* and *Vrn* genes on wheat development have been well studied. There have been detailed studies on the possibilities of various combinations of these genes under different environments to develop a tailored cultivar to achieve desired time to anthesis with different combinations of pre-anthesis phases (e.g. for *Vrn*^17,18^; for *Ppd*^12,19,20^; for combination of *Ppd* with *Vrn* or *Eps* or both^21–25^). In addition, cultivars carry residual differences in phenology after the photoperiod and vernalisation requirements have been completely satisfied, ascribed as earliness *per se* (*Eps*). In the past, the process of selecting for *Ppd* and *Vrn* genes has fixed whole breeding pools for a particular combination of photoperiod sensitivity and growth habit but *Eps* genes continue to segregate^26^. Having much smaller effect than *Ppd* and *Vrn, Eps* genes are ideal for fine-tuning wheat development^21^. The direct effect of *Eps* is known to be on heading or anthesis time, although few studies^27,25^ (*Eps-A*^*m*^ *1-l* of T. monococcum) have shown effects of *Eps* on development of spikelets. Therefore, it can be speculated that the indirect effect of *Eps* on yield could be the reason for their indirect selection in the present cultivars^25^. Current cultivars may have a reasonable variation of *Eps* alleles, providing opportunities to study these alleles in order to improve our understanding on their effects on phenology as well as on yield.

While so much is known about wheat phenology as affected by *Ppd* and *Vrn* genes, there have been fewer studies on effects of *Eps* genes on time to anthesis, and even less on whether and how the rate of organ initiation is affected. These are areas that demand attention as *Eps* genes are quite numerous (virtually across the whole genome of wheat^21,28,29^), and each potentially having a different mode of action (beyond that all *Eps* produce relatively minor changes in time to anthesis). Dissecting the effects of *Eps* into vegetative or/and reproductive phases could be crucial when considering likely effects on yield^30^, more so on how development of organs are altered. The reports on effects of *Eps* have been inconsistent so far as some studies have reported effects limited to earlier phases^25,27,31,32^ and others to that of LRP^33,34^. These studies have shown context dependent expression of traits to the extent that the same allele has differential responses based on the background it is introgressed into^35^ and epistatic interaction with respect to *Ppd* and *Vr*n genetic factors^23,25^ along with different magnitude of interaction with the growing temperature^33^. Furthermore, as all different *Eps* genes may affect development differently, it may also be possible that they might interact in determining the effects on developmental traits. To the best of our knowledge there have been studies comparing in the same experiments different *Eps* genes^35,36^, but the interaction between different *Eps* genes has never been analysed (i.e. whether the effect of a particular *Eps* gene is affected by presence of another *Eps*), which is important as many *Eps* genes are acting simultaneously in any genotype.

In the present study we aimed to analyse for the first time different developmental traits as affected by two newly identified *Eps* QTLs on chromosomes 7D and 2B and also their interactions (i.e. to what degree the effects of each of them depend upon the allelic form of the other). For that purpose, we evaluated the effect of *Eps-7D* on contrasting *Eps-2B* backgrounds and *vice–versa* on (i) phenology, not only determining time to anthesis but also quantifying whether they mainly affect development before or after TS; (ii) rate of leaf appearance; (iii) dynamics of leaf, spikelet and floret primordia development; and (iv) spike fertility.

## Results

### Heading date QTL

Three heading date QTL were identified in the Paragon x Baj RIL population with genomic locations on chromosomes *2B, 2D* and *7D* (see Table 1 and Supplementary Fig. S1). In all cases early alleles were carried by Baj. The *2D* QTL corresponds to the location of *Ppd-D1* and Baj carries the common *Ppd-D1a* allele typical of most CIMMYT varieties. The *2B* QTL is not *Ppd-B1*. On *7D* we identified a major QTL with similar additive effect to *Ppd-D1* (under UK conditions). Although specific tests of the genotypic differences in time to heading under saturating long days were not made, we assumed in principle that these genetic factors identified in chromosomes *2B* and *7D* are “earliness per se” (*Eps*) QTLs, simply based on the fact that they are definitively not *Ppd* genes and still produced significant differences in time to heading in a spring wheat background.

**Table 1.** Summary of the QTL result for heading date in the UK environment. Abbreviations: chr = chromosome, %var = percentage variance explained by the QTL, start and end marker border the QTL confidence interval.

| chr | LOD | %var | add eff | peak marker | start marker | end marker | Increase allele |
| --- | --- | --- | --- | --- | --- | --- | --- |
| **2B** | 4.3 | 6.0 | -2.12 | AX-94940971 | AX-94408000 | AX-94940971 | Baj |
| **2D** | 16.4 | 27.0 | -3.87 | AX-94603120 | AX-95217264 | AX-95124335 | Baj |
| **7D** | 10.7 | 16.2 | -3.25 | AX-94545759 | AX-94935560 | AX-94523269 | Baj |

### Phenology

Despite that these two genes had been characterised as *Eps* factors modifying time to anthesis, in the present study only the *Eps-7D* affected this trait consistently, with a large direct effect (Table 2). The direct effect of *Eps-2B* was not significant; but there was interaction between the two loci (Table 2). This means that the magnitude of the effect on time to anthesis of these *Eps* was dependent upon the allelic status of the other *Eps*. Thus, although duration from sowing to anthesis was affected by *Eps-7D* always (i.e. in both seasons and with both *Eps-2B* alleles), the magnitude of effect was different depending on the *Eps-2B* allele in the background (Fig. 1a,b). The presence of the *Eps-7D*-*early* allele shortened the time to anthesis compared with *Eps-7D*-*late* by c. 90 ºC d consistently across the two growing seasons when the background had the *late* allele of *Eps-2B* (Fig. 1a). Whereas when the background had the *Eps-2B-early* allele, the presence of the *Eps-7D*-*early* advanced anthesis only c. 45 ºC d in comparison with the *Eps-7D*-*late*, the difference was in general smaller yet significant in both seasons (Fig. 1b). Regarding the *Eps-2B* the difference in time to anthesis between lines having the early and late allele was only significant in the first season if the *Eps-7D* in the background was the *late* allele (in the second season the difference was in the same direction but only a trend; Fig. 1c), with no difference at all between the lines with contrasting *Eps-2B* alleles when the *Eps-7D-early* was in the background (Fig. 1d).

**Table 2.** ANOVA for time to anthesis. Factors that significantly affected this trait are in bold.

| Source of variation | Degrees of freedom | Mean squares (°C d) | F-ratio | Significance (P-value) |
| --- | --- | --- | --- | --- |
| Season | 1 | 422.73 | 1.71 | 0.212 |
| ***Eps-7D*** | **1** | **28698.98** | **116.38** | **<0.001** |
| *Eps-2B* | 1 | 478.60 | 1.94 | 0.185 |
| ***Eps-7D* x *Eps-2B*** | **1** | **2721.61** | **11.04** | **0.005** |
| Season x *Eps-7D* | 1 | 88.65 | 0.36 | 0.558 |
| Season x *Eps-2B* | 1 | 489.38 | 1.98 | 0.181 |
| Season x *Eps-7D* x *Eps-2B* | 1 | 614.34 | 2.49 | 0.137 |
| Blocks | 2 | 858.61 | 3.48 | 0.059 |
| Error | 14 | 246.61 |  |  |

**Figure 1.**
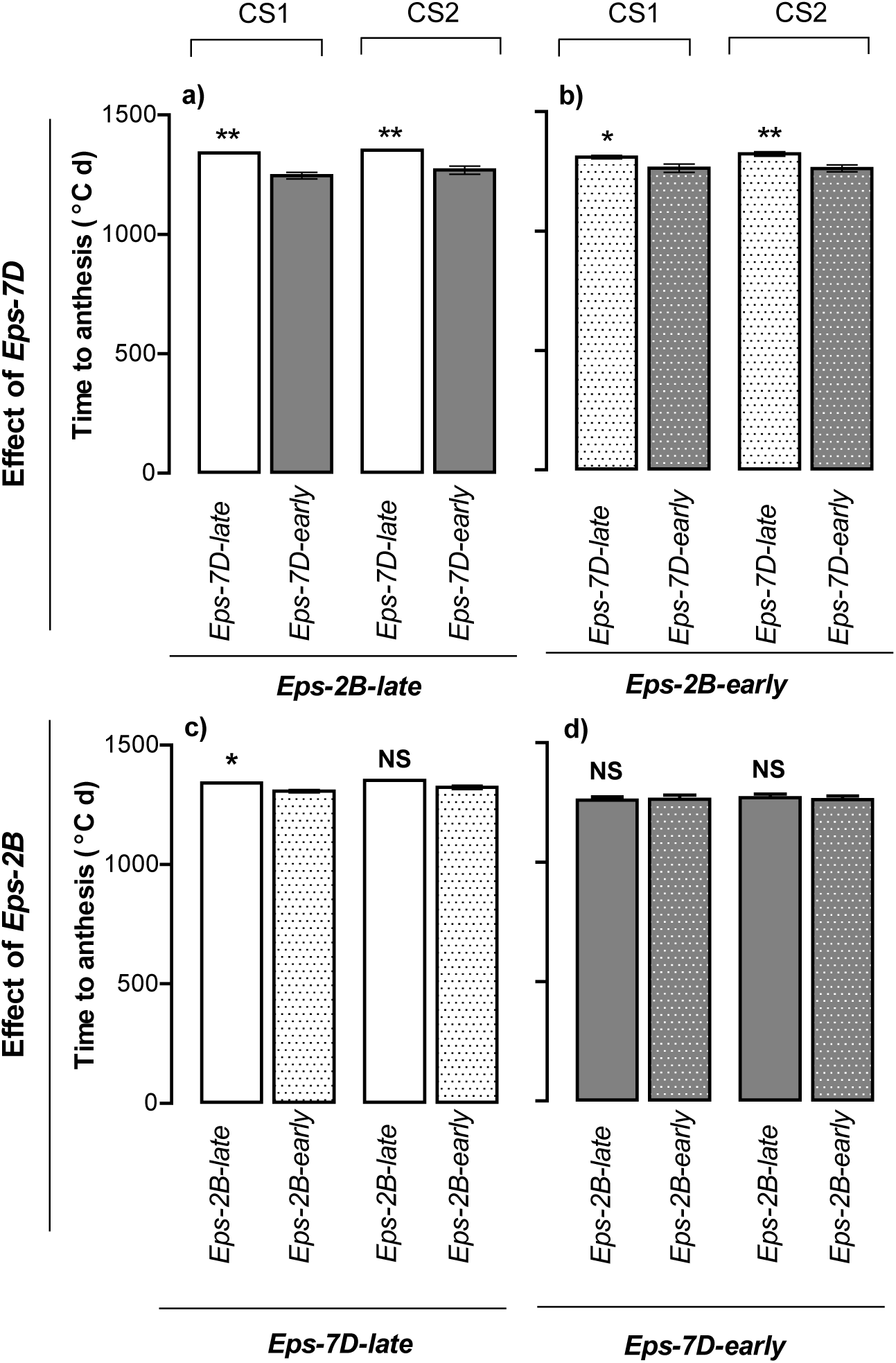
Duration of the crop cycle from sowing to anthesis for the lines carrying (i) the *Eps-7D-late* or *early* allele (upper panels) on the contrasting background of *Eps-2B* (*Eps-2B-late* and -*early*, upper panel left and right, respectively); or (ii) the *Eps-2B-late* or *early* allele (bottom panels) on the contrasting background of *Eps-7D* (*Eps-7D-late* and -*early*, bottom panel left and right, respectively). Data are shown for each of the two cropping seasons (CS1 and CS2 within each panel). Significance level shown for the differences between alleles within each season: *p<0.05; **p<0.01; NS=non-significant.

When analysing the effect of *Eps-7D* on particular pre-anthesis phases (before and after TS) the results again provided evidence for an interaction with the allelic form of *Eps-2B* gene in the background (Fig. 2). The effect of *Eps-7D* strongly and consistently affected the duration of the period until TS (c. 60 °C d averaging across the seasons; p<0.001) and only marginally affected the LRP in the second season when the *Eps-2B* gene in the background was the *late* allele (Fig. 2a,b). On the other hand, not only the overall effect of *Eps-7D* was smaller when in the background the *Eps-2B* was the *early* allele (Fig. 1c) but also it was mainly due to its effect on the LRP (c. 44 °C d averaging across the seasons; p<0.001) with no changes in time to TS at all (Fig. 2e,f). The *Eps-2B* gene only affected the measured phenology traits when the *Eps-7D* in the background was the *late* allele (Fig. 2c,d), with non-significant differences when the *Eps-7D-early* allele was in the background (Fig. 2g,h). In that case and consistently across both seasons, the effect was seen in the early development up to TS (c. 52 °C d averaging across the seasons; p=0.08) with no clear effects on the duration of LRP (Fig. 2c,d).

**Figure 2.**
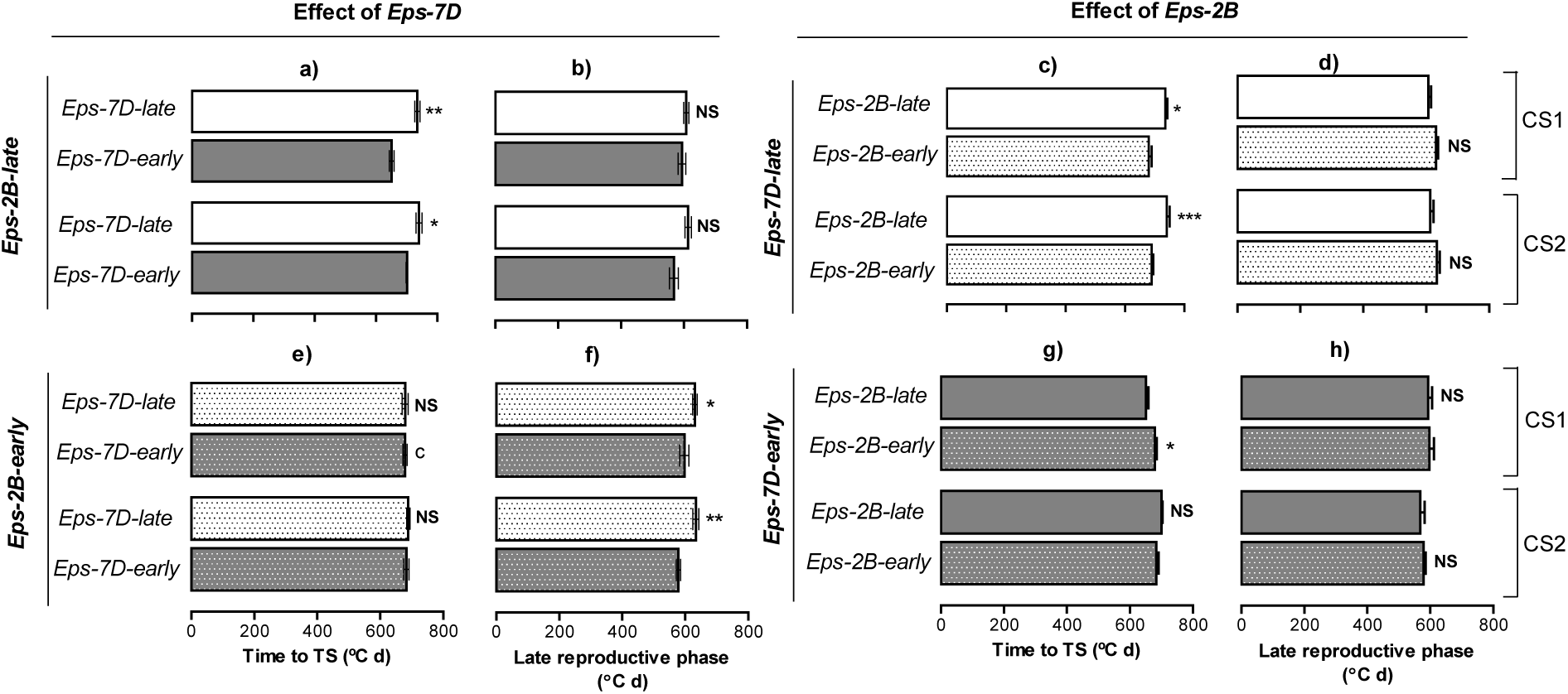
Duration of the two pre-anthesis phases considered: time from sowing to terminal spikelet (TS) and from then to anthesis, the late reproductive phase as affected by *Eps-7D* (left panels) and *Eps-2B* genes (right panels) on backgrounds contrasting in the allelic form of the other *Eps* gene (top and bottom panels for the late and early alleles of the other *Eps* gene). Data are shown for each of the two cropping seasons (CS1 and CS2). Segments in each bar stand for the SEM. Significance level shown for the differences between alleles within each season: *p<0.05; **p<0.01; NS=non-significant.

### Dynamics of leaf appearance and final leaf number

There was no significant effect of the *Eps-7D* gene on the final leaf number (FLN), lines with the *Eps-7D-late* and -*early* alleles produced 9.55 and 9.57 leaves (averaged across the two *Eps-2B* alleles in the background and seasons), respectively (Fig. 3a,b,e,f). On the contrary presence of the allele *Eps-2B-early* tended to reduce the FLN (Fig. 3c,d,g,h). But this effect exhibited an interaction with the allelic form of the *Eps-7D* gene in the background. Lines with *Eps-2B-early* showed c. 0.7 leaves less than *Eps-2B-late* lines (averaged across seasons) when the background was the *Eps-7D-late* (Fig. 3c,g), while the difference was less clear (c. 0.4 leaves), and statistically not significant, when in the background the *Eps-7D* gene had the early allele (Fig. 3d,h).

**Figure 3.**
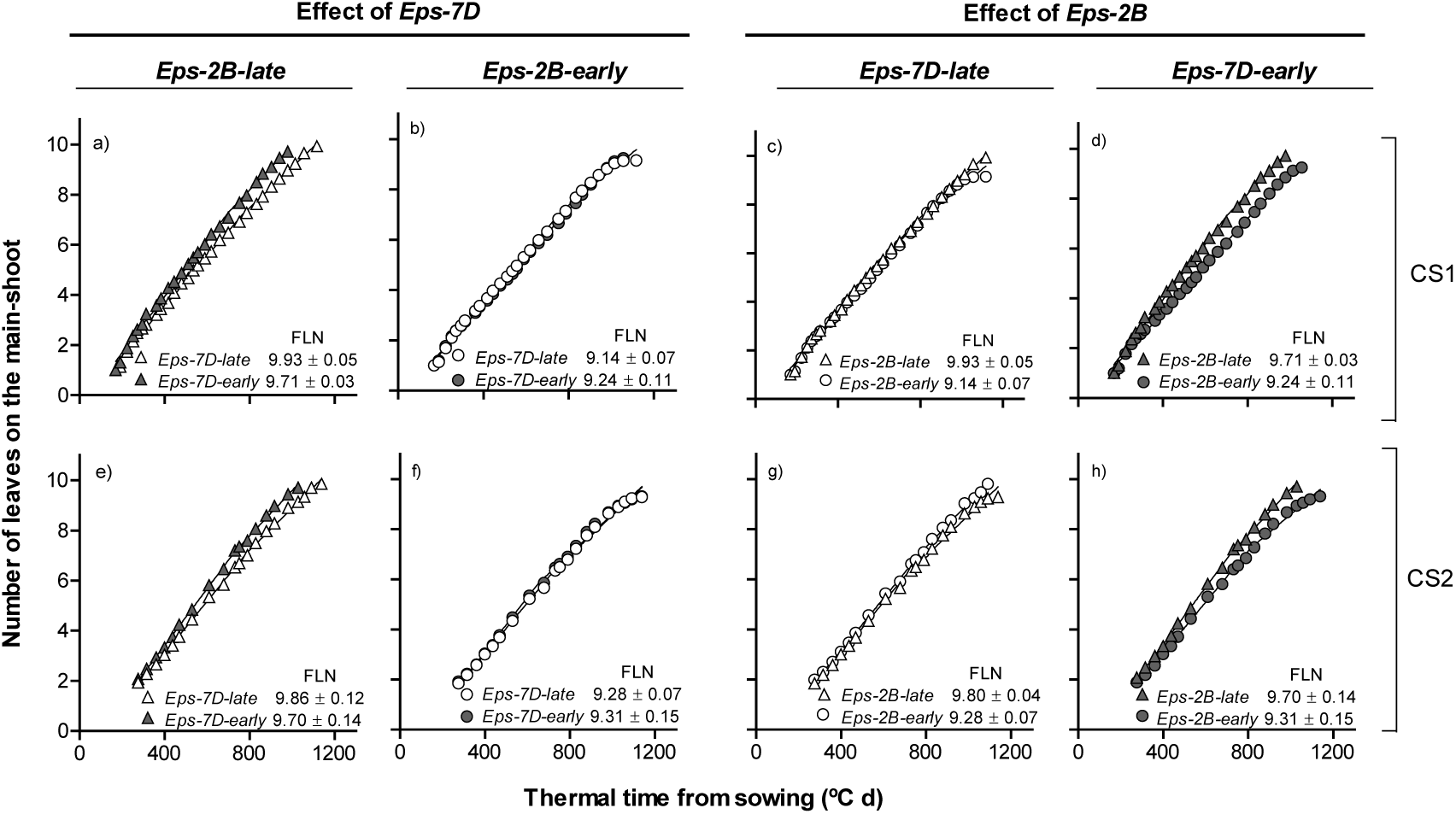
Relationship between number of leaves appeared on the main shoot and thermal time from sowing as affected by *Eps-7D* (left panels: a, b, e and f) and *Eps-2B* genes (right panels: c, d, g and h) on backgrounds contrasting in the allelic form of the other *Eps* gene (left and right panels within each *Eps* gene) in cropping seasons 1 (top panels: a-d) and 2 (bottom panels: e-h). Inside each of the panels are the final leaf number (FLN) with their SEM.

Although in all cases the relationship between the number of appeared leaves and thermal time had a strong linear component (Fig. 3), the relationships were actually bi-linear with early leaves appearing significantly faster (and having therefore a shorter phyllochron) than later leaves (Table 3). Opposite to the differences between *Eps* genes on FLN, the *Eps-7D* gene affected phyllochron (in general both that of the early and late leaves and regardless of the *Eps-2B* allele in the background) and in line with the effects of this gene on time to anthesis the effect was smaller and less clear in the second growing season (Table 3, top half). The effects of *Eps-2B* on phyllochron was not clear, as differences between lines with the *Eps-2B*-*early* and *Eps-2B*-*late* were small and inconsistent (Table 3, bottom half).

**Table 3.**
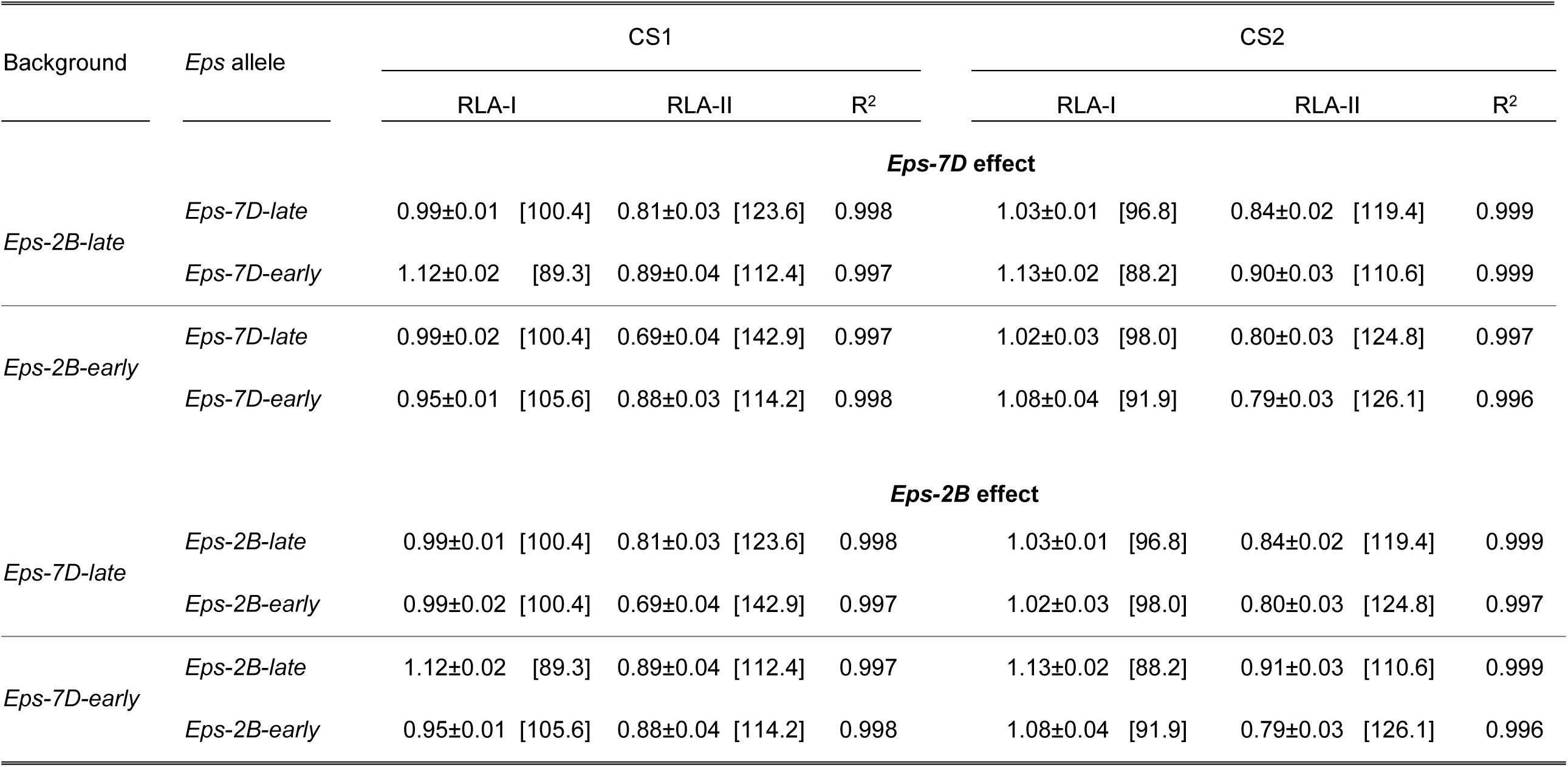
Rates of leaf appearance (leaves [100 ºC d]^-1^; ±SE) corresponding to the early- and late-appearing leaves (RLA-I and RLA-II, respectively) as affected by *Eps-7D* and *Eps-2B* genes (top and bottom parts of the Table, respectively) on backgrounds contrasting in the allelic form of the other *Eps* gene in the two cropping seasons (CS1 and CS2), and coefficients of determination of the segmented linear regression (in all cases P<0.001). The corresponding phyllochron values (ºC d leaf^-1^) are included between square brackets.

### Dynamics of leaf and spikelet primordia development

The dynamics of leaf and spikelet primordia initiation showed that, in general, both *Eps-7D* and *Eps-2B* effects increased the number of primordia initiated when lines carrying the *late* allele were compared with those with the *early* allele, particularly when the other *Eps* gene in the background was the *late* allele (Fig. 4). As differences in FLN were small or negligible (see above), these differences in number of primordia reflected those in spikelets initiated per spike. Overall, none of the two *Eps* genes affected the rate of primordia initiation in any of the two cropping seasons (Fig. 4). Thus, the differences in the total number of primordia initiated between lines carrying the *late* and *early Eps* alleles were mostly due to the differences in duration of primordia initiation (Fig. 4).

**Figure 4.**
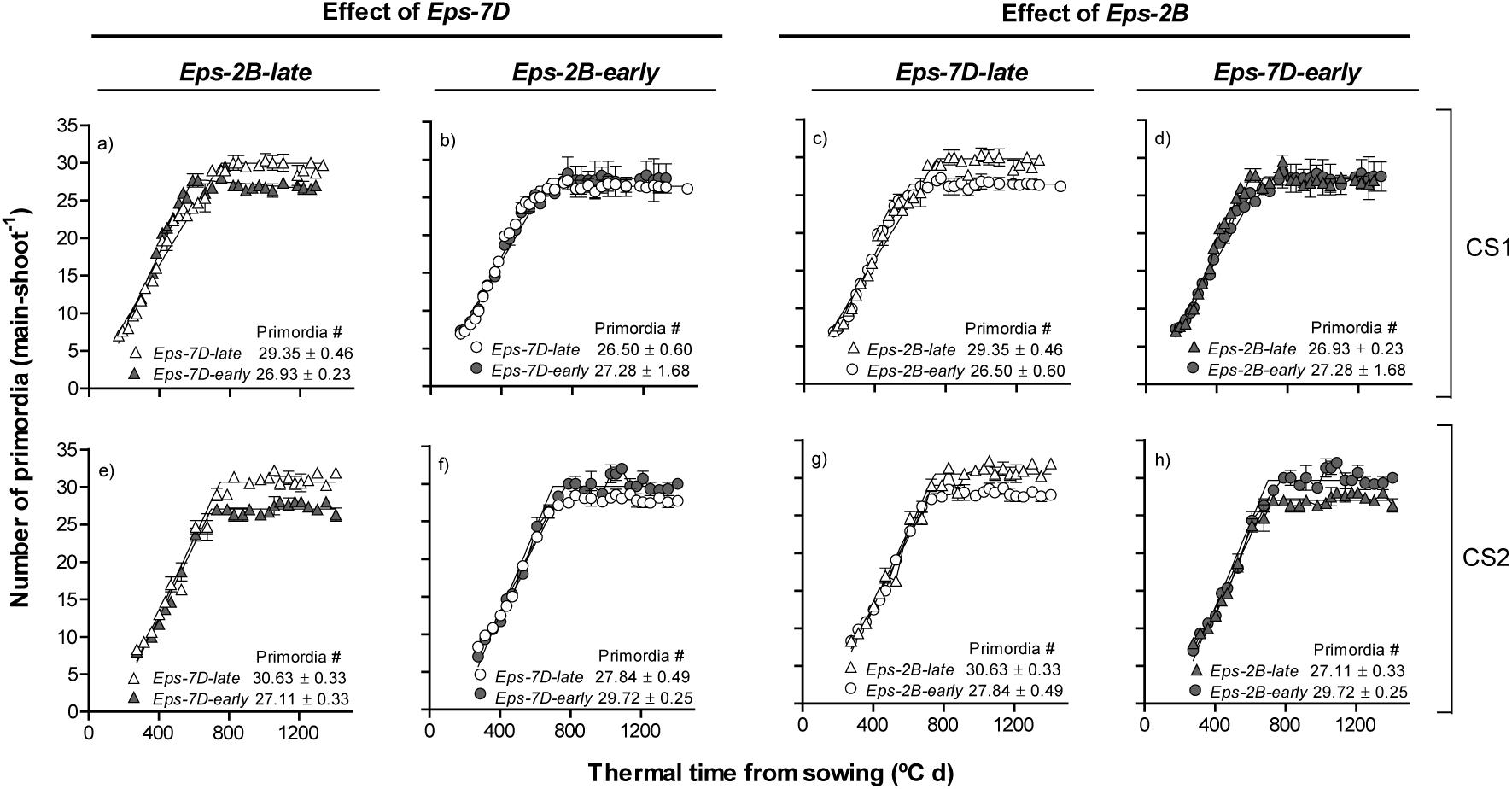
Relationship between number primordia initiated on the main shoot apex and thermal time from sowing as affected by *Eps-7D* (left panels: a, b, e and f) and *Eps-2B* genes (right panels: e, d, g and h) on backgrounds contrasting in the allelic form of the other *Eps* gene (left and right panels within each *Eps* gene) in cropping seasons 1 (top panels: a-d) and 2 (bottom panels: e-h). Inside each of the panels are the total number of primordia with their SEM.

### Dynamics of floret development

Detailed study of developmental dynamics of each individual floret initiated in 3 spikelet positions of the spike (apical, central and basal spikelets) was carried out to understand the basis of *Eps-7D* and *Eps-2B* spike fertility effects. In order to focus on aspects of floret dynamics which directly influences grain number we only presented the dynamics of florets that reached at least the stage W4.5 (carpel primordia visible); i.e. up to the fifth floret from the rachis. Outside of this range seeds are hardly ever produced. At the other end of the floret survival scale the two florets most proximal to the rachis (F1 and F2) are very similar and the likelihood of survival was always 100% for these florets, hence we focused here on the developmental progress of the F2, F3, F4 and F5 florets to reflect any effects of *Eps* genes on floret fertility.

For the same reason, as results were consistent across growing seasons, we only included data for the first season in the main text and sent the equivalent figures for the second season to supplementary materials. Regarding the effect of *Eps-7D* on development of florets, the interaction with the allele of *Eps-2B* in the background was evident as well when comparing the differences between lines with *Eps-7D*-*early* and *Eps-7D*-*late* between when in the background the *Eps-2B* was the *late* or *early* allele. When the background had the *Eps-2B*-*late* development of individual floret primordia was in general faster in lines with the *Eps-7D*-*early* allele and this resulted in a different likelihood of some labile florets to become fertile at anthesis (Fig. 5 and Supplementary Fig. S2). There was no difference in final fertility of the F2 floret among any of the lines in any of the three spikelet positions in both the seasons: F2 always reached the fertile floret stage (Fig. 5a,e,i and Supplementary Fig. S2a,e,i). However, and even when not producing a difference in fertility for this particular proximal floret, the rate of development seemed consistently faster for this F2 for lines with *Eps-7D*-*early* allele (Fig. 5a,e,i and Supplementary Fig. S2a,e,i). The remaining distal florets (F3, F4 and F5) also showed the same trend of faster rate of floret development in lines carrying *Eps-7D*-*early* than those carrying *Eps-7D*–*late*, affecting the likelihood of these florets becoming fertile depending on the particular floret and spikelet positions. Despite the different rates of floret development, F3 reached similar final levels of fertility in central spikelets in lines with any of the alleles of *Eps-7D*; but in the basal and apical spikelets the lower rate of floret development in lines with *Eps-7D*-*late* allele determined that F3 was fertile in only c. 33% of the plants measured whilst it was fertile in almost all plants of the lines with the *Eps*-7D-*early* allele (Fig. 5b,j and Supplementary Fig. S2b,j). When considering F4 these lines differed in the time of initiation along with rate of development and plants with *Eps-7D*-*early* had earlier initiation and faster rate of development of F4 than those with *Eps-7D*-*late*. When the *Eps*-7D was the *early* type c. 60% of F4 reached W10 in apical and basal position, while they were never fertile in lines with the *Eps-7D*-*late* allele (Fig. 5c,k and Supplementary Fig. S2c,k). Floret F5 reached W4.5 in almost all the lines but was fertile only in the central spikelet of most, but not all, plants with the *Eps-7D*-*early* (Fig. 5h and Supplementary Fig. S2h). When we compared the rates of developments and fate of floret primordia of lines with the *Eps-7D*-*early* and –*late* alleles but with the *Eps-2B*-*early* in the background, there were no noticeable differences in any of the spikelets and floret positions (Fig. 6 and Supplementary Fig. S3). That is, the improved rates of development in the lines with the *Eps-7D*-*early* allele evidenced consistently across floret and spikelet positions and over the two growing seasons when the background was the *Eps-2B-late* were not evident anymore. Regarding the effect of *Eps-2B* on development of florets, the effects were more clear when in the background the *Eps-7D* was the *early* allele and generally subtler than those of *Eps-7D* (Fig. 7, 8 and Supplementary Fig. S4, S5). When the background had the *Eps-7D*-*late* development of individual floret primordia was rather similar in all lines regardless of the *Eps-2B* allele (Supplementary Fig. S4). On the other hand, when the *Eps-7D* in the background was the *early* allele, there were no clear differences in rates of development for most proximal florets but for the labile floret primordia (i.e. those reaching W10 or dying depending on the conditions) lines with the *Eps-2B*-*late* allele had maintained a faster rate of development than in lines with the *Eps-2B-early* allele, allowing these labile florets reaching the W10 stage in the former, (as well as attaining higher floret score in florets not being fertile in any of the lines; Fig. 8 and Supplementary Fig. S5).

**Figure 5.**
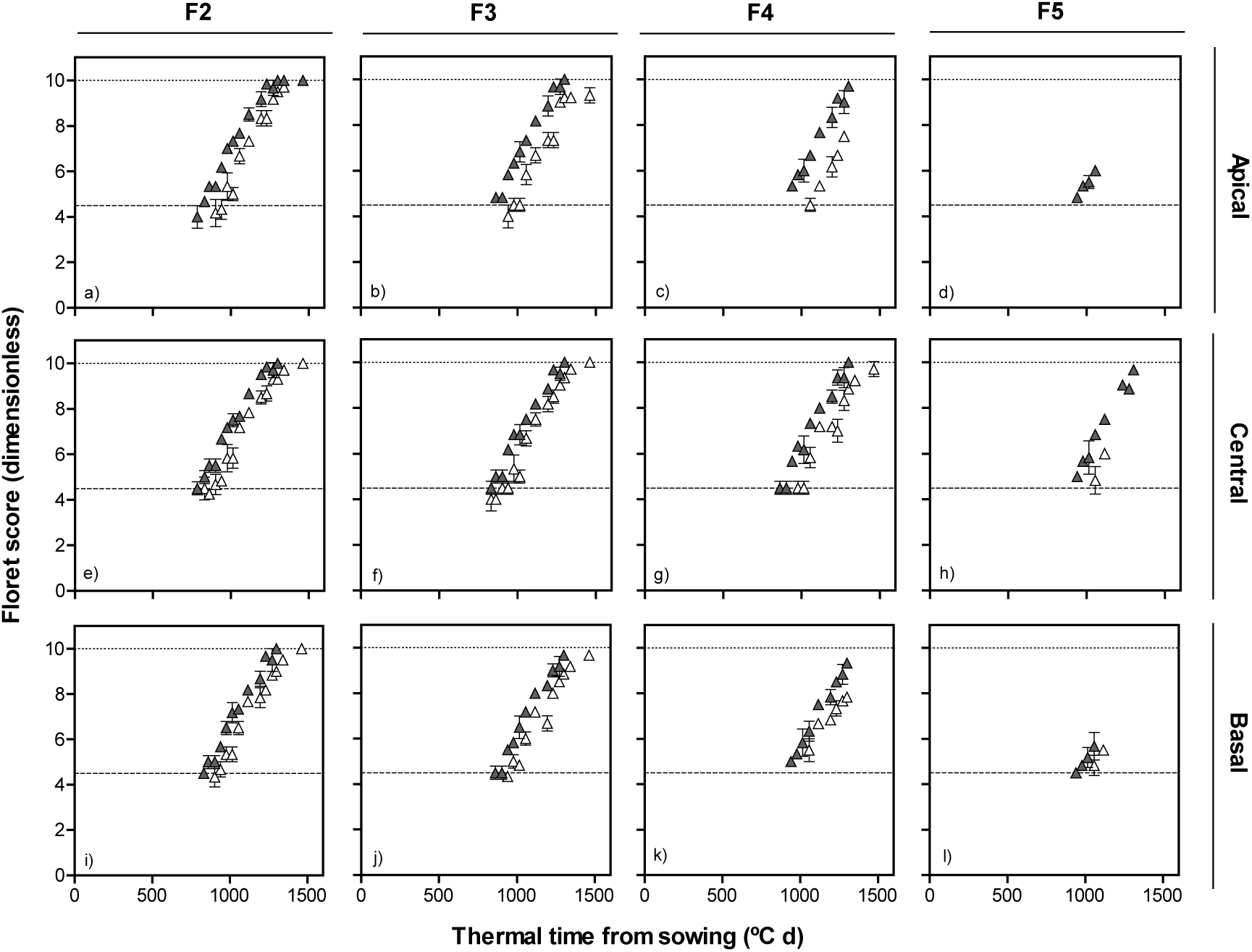
Dynamics of floret development (floret score) in F2, F3, F4 and F5 florets at apical (top panels: a-d), central (middle panels: e-h) and basal (bottom panels: i-l) positions of the spike with thermal time from sowing in lines with *Eps-7D-late* (open symbol) and -*early* (closed symbol) allele with *Eps-2B-late* allele in the background in the first cropping seasons.

**Figure 6.**
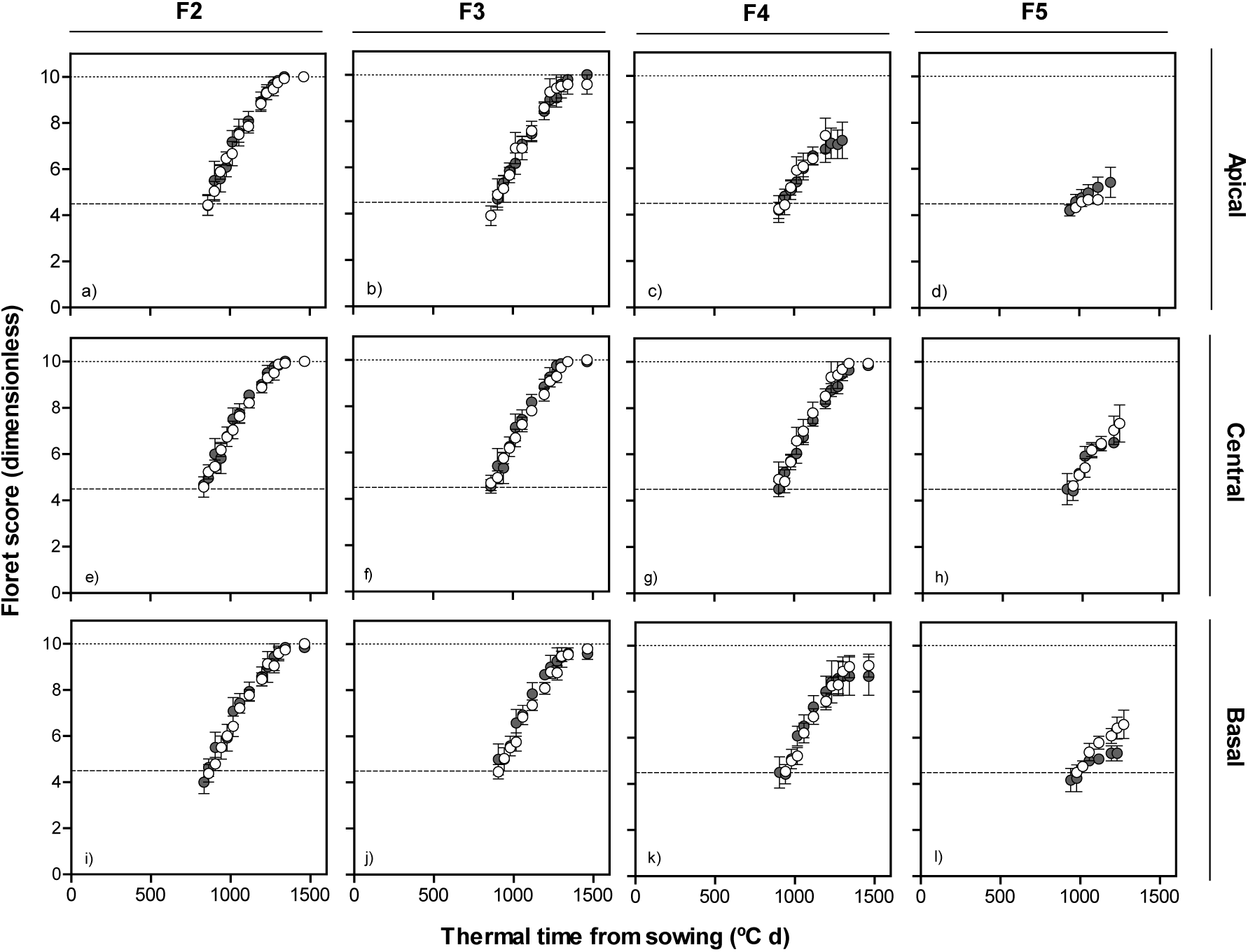
Dynamics of floret development (floret score) in F2, F3, F4 and F5 florets at apical (top panels: a-d), central (middle panels: e-h) and basal (bottom panels: i-l) positions of the spike with thermal time from sowing in lines with *Eps-7D-late* (open symbol) and *early* (closed symbol) allele with *Eps-2B-early* allele in the background in the first cropping season.

**Figure 7.**
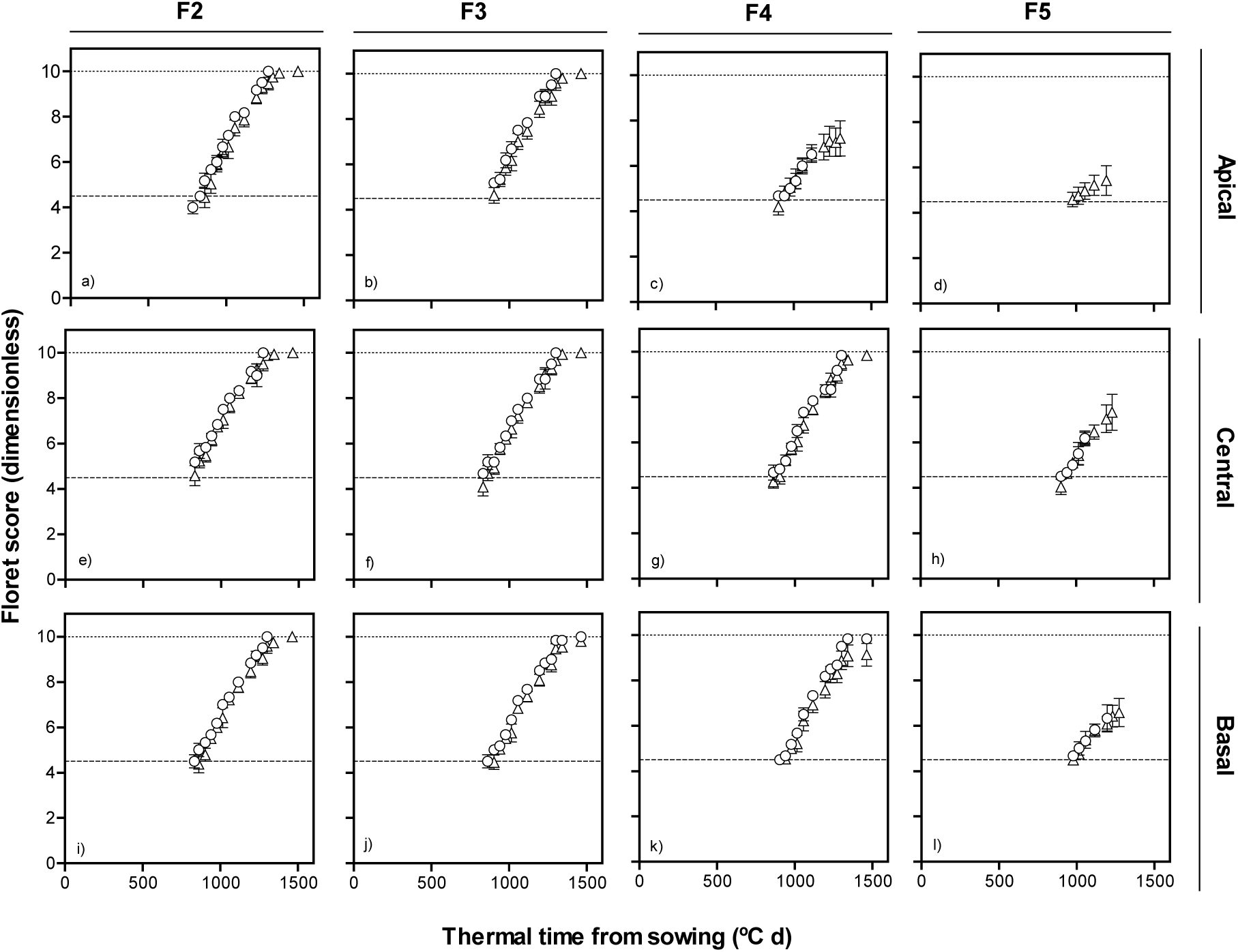
Dynamics of floret development (floret score) in F2, F3, F4 and F5 florets at apical (top panels: a-d), central (middle panels: e-h) and basal (bottom panels: i-l) positions of the spike with thermal time from sowing in lines with *Eps-2B-late* (triangles) and *early* (circles) allele with *Eps-7D-late* allele in the background in first cropping season.

**Figure 8.**
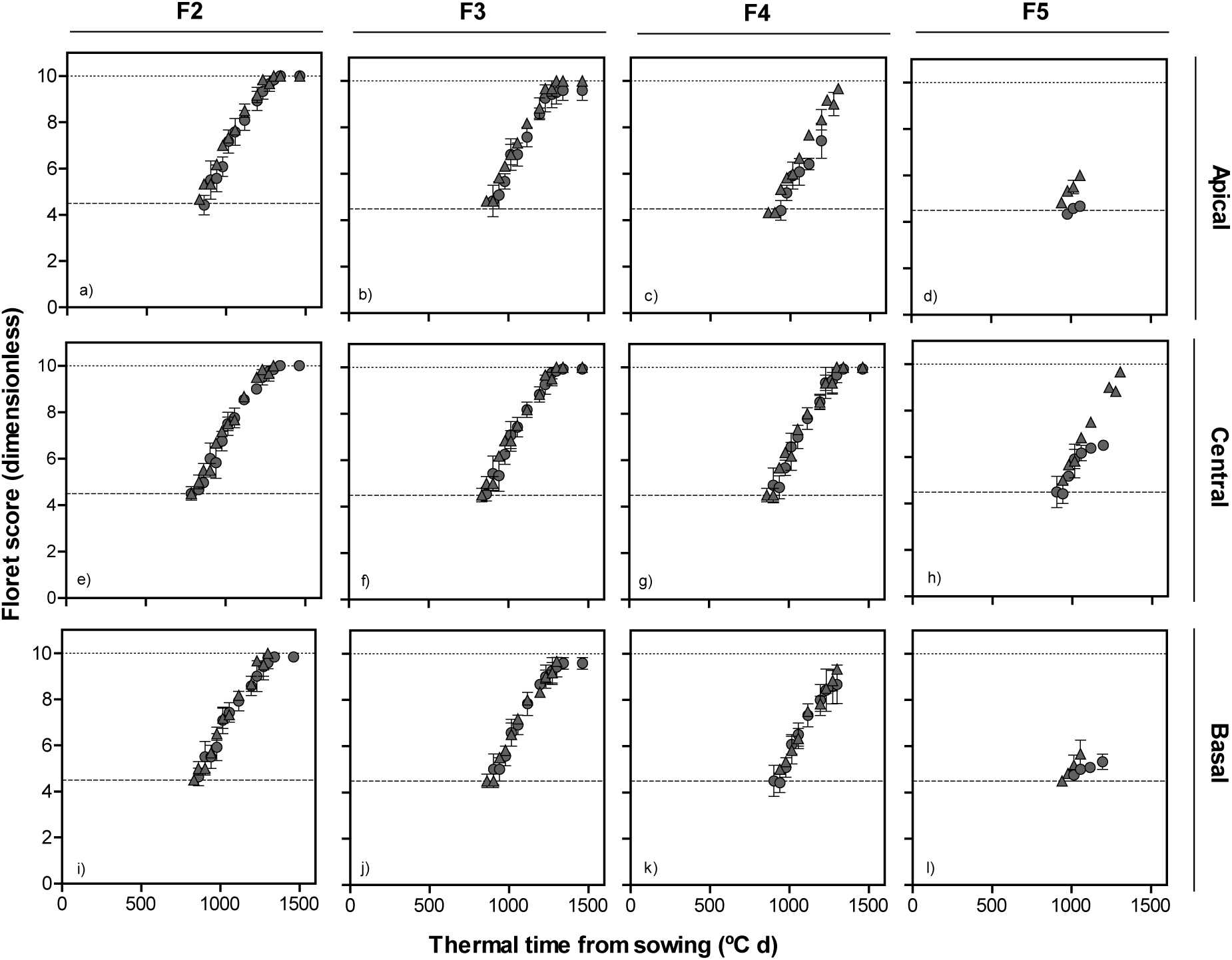
Dynamics of floret development (floret score) in F2, F3, F4 and F5 florets at apical (top panels: a-d), central (middle panels: e-h) and basal (bottom panels: i-l) positions of spike with thermal time from sowing in lines with *Eps-2B-late* (triangles) and -*early* (circles) allele with *Eps-7D-early* allele in the background in first cropping season.

### Floret primordia initiation and death

Detailed analysis was carried out to calculate the living floret primordia (again considering only floret primordia that reached a score ≥ W4.5, and therefore the maximum number of primordia was “only” 6-7). The patterns of floret initiation and mortality for each genotype were relatively similar for the three different spikelet positions considered (Supplementary Fig. S6 and S7) and therefore we reported here an average of these three spikelets (Fig. 9). The largest effect was that produced by the *Eps-7D* gene when in the background the allele of the other *Eps* gene was *Eps-2B*-*late*. In this case the lines carrying the *Eps-7D*-*early* produced more floret primordia (reaching W4.5 or higher; if all florets visible microscopically were taken into account there would be no difference in maximum number of floret primordia) and increased the rate of floret survival, both contributing to an improved number of fertile florets compared with line carrying the *Eps-7D*-*late* (Fig. 9a). These *Eps*-7D-*early* lines initiated floret development earlier and therefore the duration of the process of floret initiation was not reduced (Fig. 9a), although the duration of the LRP tended to be shorter (Fig. 2b). When the *Eps-2B* in the background was the *early* allele the *Eps-7D* showed no clear effect on the dynamics of floret initiation-mortality (Fig. 9b). Considering the effect of *Eps-2B*, there was no clear trends in the dynamics of floret initiation/mortality when the *Eps-7D* in the background was the *late* allele (Fig.9c). However, when the background had the *Eps-7D*-*early* the *Eps-2B*-*late* slightly increased spike fertility with respect to *Eps-2B*-*early* mainly through improving floret primordia survival, as the maximum number of floret primordia initiated (and reaching at least W4.5) was similar (Fig. 9d).

**Figure 9.**
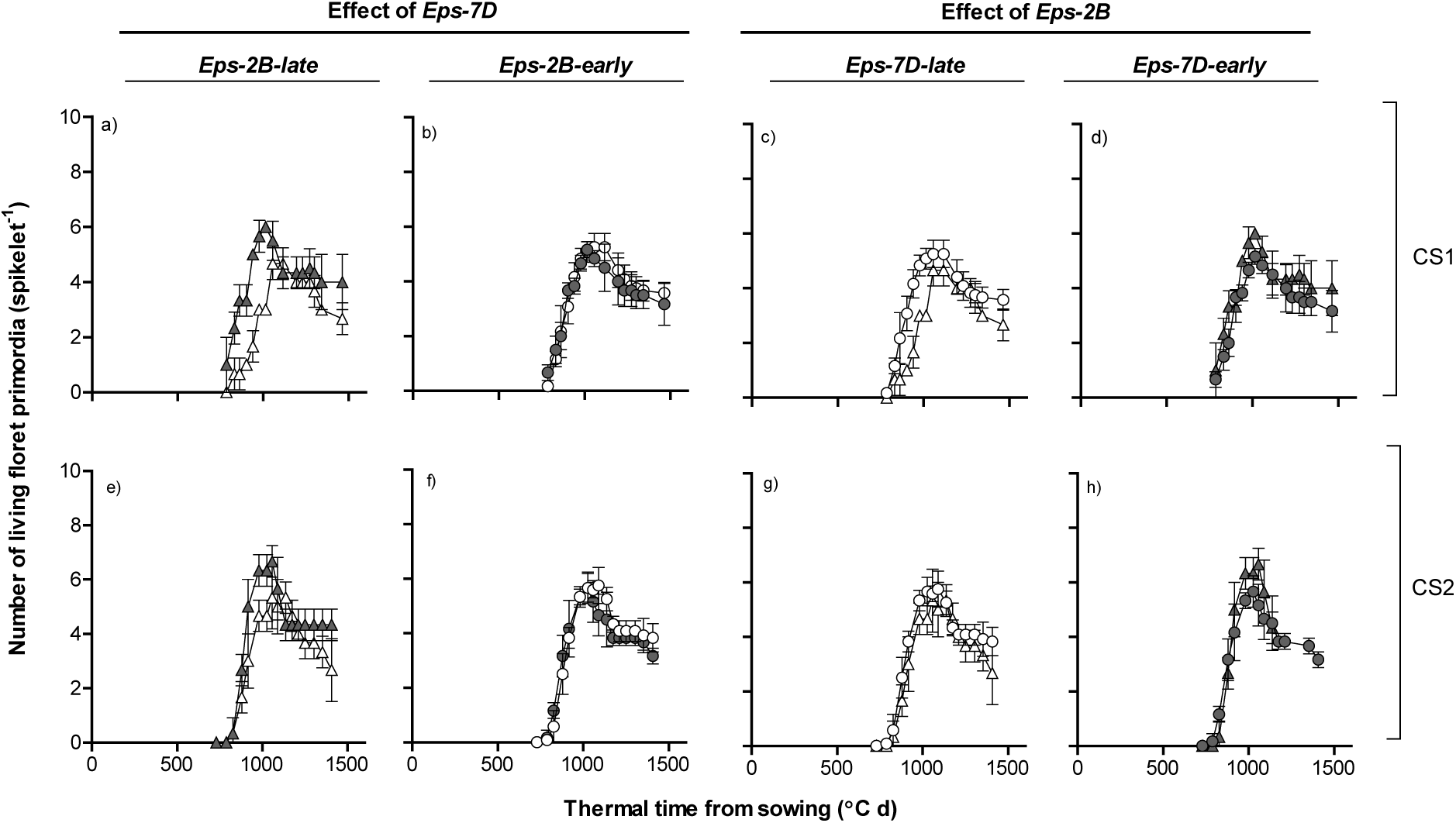
Number living floret primordia per spike and thermal time from sowing as affected by *Eps-7D* (left panels: a, b, e and f) and *Eps-2B* genes (right panels: c, d, g and h) on backgrounds contrasting in the allelic form of the other *Eps* gene (left and right panels within each *Eps* gene) in first (top panels: a-d) and second (bottom panels: e-h) cropping seasons. *Eps-7D-late* and -*early* (open and closed symbols, respectively) and *Eps-2B-late* and -*early* allele (triangle and circles, respectively).

### Fertile florets at anthesis

There was a clear interaction between *Eps-7D* and *Eps-2B* on the number of fertile florets per spike. Lines carrying the *Eps-7D*-*early* allele had higher spike fertility than those with the *Eps-7D*-*late* alleles when the other gene in the background was *Eps-2B*-*late* allele (Fig. 10a,e), but not when the *Eps-2B* had the *-early* allele (Fig. 11b,f). Reflecting the interaction between the two *Eps* genes, the *Eps-2B* also affected significantly spike fertility when the *Eps-7D* in the background was the *early* type (Fig. 10d,h), and the magnitude of the difference shrunk when the *Eps7D*-*late* was in the background (Fig. 10c,g). Differences in fertile florets per spike between lines with contrasting forms of the *Eps-7D* were concentrated in central spikelets of the spike and the advantage was substantial enough to override the fact that the lines with *Eps-7D*-*late* produced more spikelets per spike (Fig. 10a,e). On the other hand, the effect of *Eps-2B* on fertile florets was subtle and not significant yet very consistent compared to that of *Eps-7D*, and differences in floret fertility were relatively minor but consistent across most spikelets (Fig. 10d,h).

**Figure 10.**
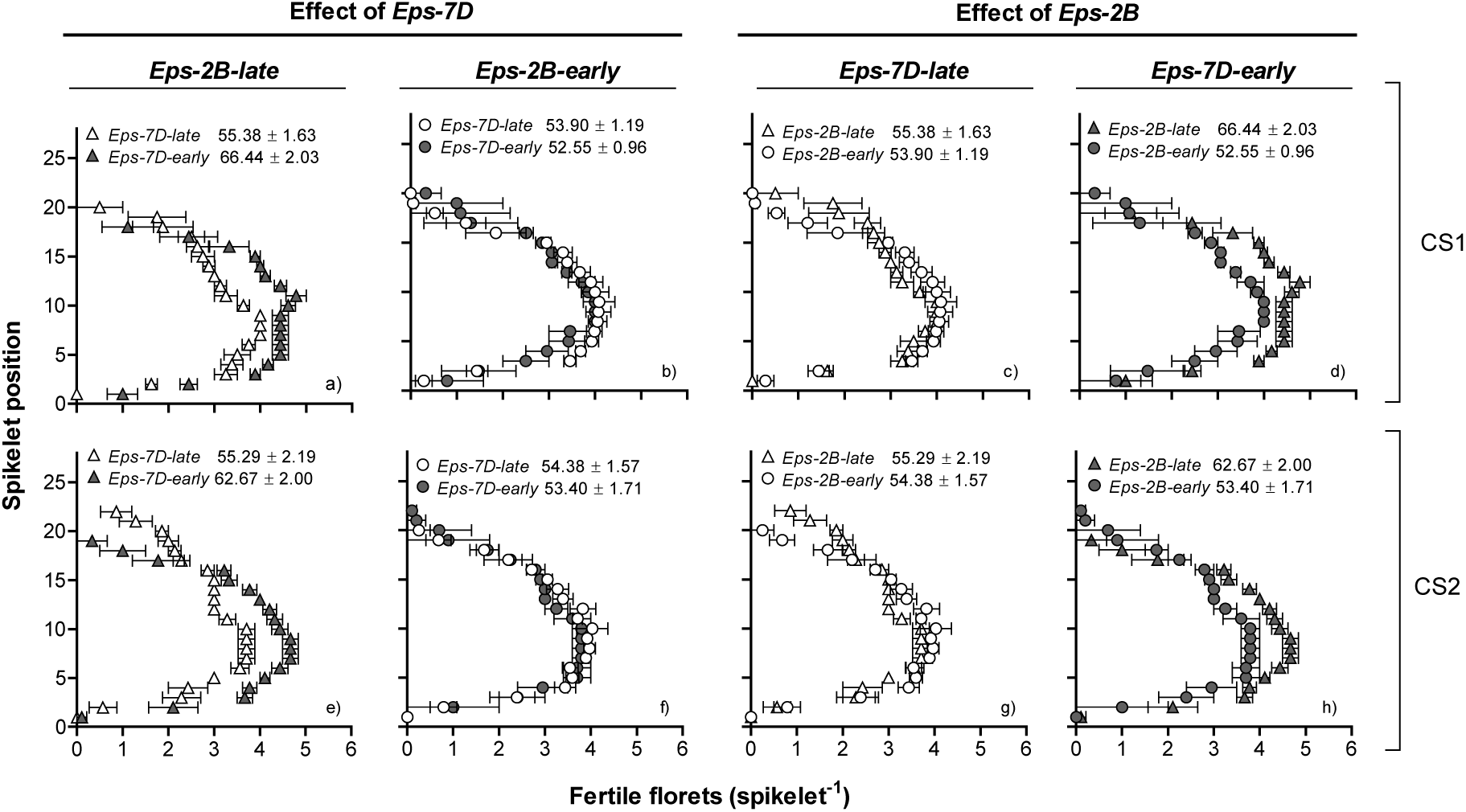
Mapping of fertile florets at anthesis per each spikelet in the spike as affected by *Eps-7D* (left panels: a, b, e and f) and *Eps-2B* genes (right panels: c, d, g and h) on backgrounds contrasting in the allelic form of the other *Eps* gene (left and right panels within each *Eps* gene) in first (top panels: a-d) and second (bottom panels: e-h) cropping seasons. Inside each of the panels are the fertile florets per spike with SEMs.

**Figure 11.**
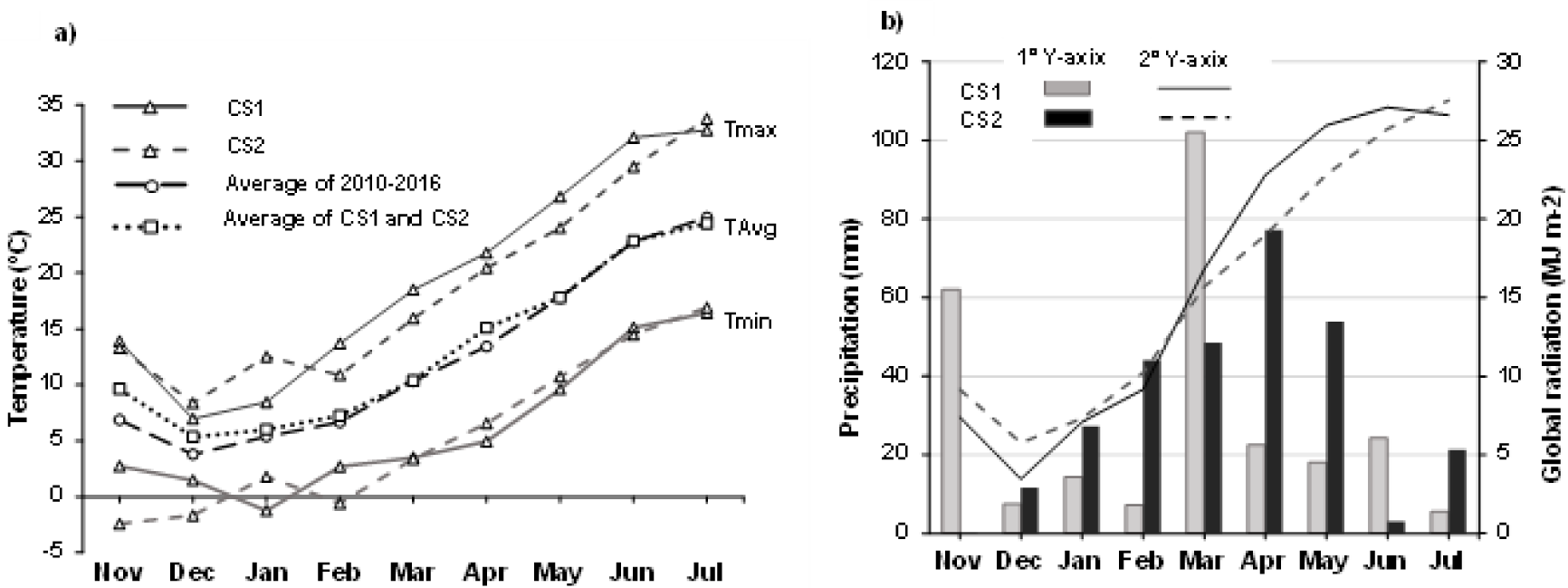
Monthly minimum (Tmin), maximum (Tmax) for first (CS1) and second (CS2) cropping season and mean of the monthly average temperature (TAvg) of CS1 and CS2 as well as monthly TAvg of cropping seasons from the past 5 years (2010-2016, a); monthly accumulated precipitation and global radiation for CS1 and CS2 (b).

## Discussion

The two newly identified *Eps* QTLs, *Eps-7D* and *2B* affected the whole cycle from sowing to anthesis but the effects were influenced by the allele of the other *Eps* in the background. The overall effect of these two *Eps* genes was relatively small, although *Eps-7D* was stronger than *Eps-2B*. The effect of *Eps-7D* on time to anthesis was as strong as it has been reported before for other *Eps* genes (e.g. ^21,24,26,27,35,37,38^) but effect of *Eps-2B* was relatively very small again such smaller effects were also noted for other *Eps*^35^. Therefore, either of them, but particularly the *Eps-7D* due to its more consistent effect, might be exploited to fine-tune time to anthesis of elite germplasm.

It is true that the changes in the whole phase from sowing to anthesis can be brought about by various combination of changes in its component pre-anthesis phases (with more or less independency) and its importance lies in the fact that each of these phases determine initiation and survival of various organs that may end up affecting yield components^39^. *Eps-7D* affected both time to TS and LRP while the subtle effect of *Eps-2B* on whole phase was only evident in time to TS^27^. Our study provides the first time proof for the interaction between two Eps in particular which was only speculated before^38^. Our observations in addition to that of other researcher ^35^ suggest that different *Eps* genes may differ in their effects on individual developmental phases, even if their effect on the overall time to anthesis were similar.

It was important to note that delay due to *late* allele of one *Eps* was enhanced in presence of the *late* allele of the other *Eps* (the line carrying *late* allele at both *Eps-7D* and *2B* had the longest duration from sowing to anthesis), pointing towards probable additive effects for the duration of developmental phases. And discrepancies about the effect of same *Eps* in the literature (under different temperatures or background) also suggests the epistatic interaction of *Eps* with other unknown flowering genes and interaction with temperature. Therefore, it cannot be overemphasized on how important it is to study individual *Eps* from a particular genetic background with known flowering gene complex (other major and minor flowering genes involved) acting in the background in order to decode the functions of *Eps* (certainly beyond time to anthesis) for exploiting them in breeding. As *Eps*, unlike *Ppd* and *Vr*n, are present in almost all of the chromosomes, their abundancy provide an enormous opportunity to work with at the same time most of them being minor genes and probable additive effect, their abundance also poses difficulty in clear understanding of their function (and pathway).

The QTLs also differed in their effect on FLN, while *Eps-7D* did not alter the FLN *Eps-2B*-*early* had significantly (though naturally only slightly) lower FLN than the respective *late* allele when the background was *Eps-7D*-*late*. Effect of *Eps-2B* on FLN support the study conducted by Hoogendoorn (1985)^40^ observing variations in FLN in their vernalized photoperiod insensitive lines. This differential effect of *Eps-7D* and *Eps-2B* on FLN when having similar effects on the duration of early phases of development shows another differential mechanisms of action of these genes in early developmental traits when seen in more detail: while both affected similarly the rate of development of the early phases (and might be presumed acted in that phase similarly), it seemed that the *Eps-7D* affected the rate of leaf initiation in parallel, and consequently the lines with the *Eps-7D*-*early* allele had the same FLN as the lines with *Eps-7D*-*late* did; while *Eps-2B* seemed to have not affected the rate of leaf initiation and therefore the effects on duration of an early developmental phase is reflected on FLN. Furthermore, the *Eps-7D* gene affected the rate of leaf appearance as the main mechanism for changing time to anthesis, as also found by Ochagavia et al. (2018)^35^ for another *Eps* gene. This opposite effects on phyllochron, are commensurate with the fact that *Eps-7D* did also affect the duration of the LRP while *Eps*-2B restricted its effect to the earlier phases.

Further, both *Eps-7D* and *2B* affected the number of primordia initiated and the differences in the primordia was mainly contributed from spikelet primordia as the differences in FLN between *late* and *early* alleles of both the QTLs were rather small. And the differences in the primordia were mainly due to the differences in the duration of primordia initiation as there was no alteration in the plastochron. Such marginal effect of *Eps* on spikelet number per spike mainly due to the *Eps* effect on phenophases without affecting the rate of primordia initiated have been observed in other studies involving ‥major‥^27^ and ‥minor‥^35,36,40^ *Eps* genes. Among the few studies reporting *Eps* effects beyond time to anthesis the studies by Lewis et al. (2008)^27^ in diploid wheat (under field condition) and Alvarez et al. (2016)^25^ in tetraploid wheat (under controlled conditions) were exceptions showing effect of *Eps-Am 1* bringing about larger changes in spikelet number per spike but this exception was justified by the unusual larger effect of that *Eps* on time to anthesis. Like other studies in the literature have reported, *Eps* QTLs bringing about significant (moderate to major) variation in developmental patterns have only brought about subtle variations in the components related to yield (spikelet number^40^;spike fertility^36^). Further, our results indicate that both the *Eps* QTLs alter developmental phase resulting in modification of anthesis time have contrasting pattern of their function on the dynamics of organs initiated.

*Eps-7D* and *Eps-2B* had contrasting effect on spike fertility presented as floret development and fertile floret per spike at anthesis, again the effect of both *Eps* on spike fertility was influenced by the presence of the *early* or *late* allele of the other *Eps* in the background. Unexpectedly, lines with the *Eps-7D*-*early* allele had a higher spike fertility compared to *late* allele despite the shorter LRP, at least when the background allele for *Eps-2B* was *late*. A general hypothesis is having a longer LRP would improve spike fertility which has a straight forward explanation that longer duration allows distal florets to become fertile along with positive effect from improved spike dry weight (SDW)^13^. But this is true only when the genetics involved in altering the duration does not affect the rate of development and the differences in the duration was big enough to cause variations in SDW. In this particular case, the situation was different as the period of floret development was similar between lines with contrasting alleles of *Eps-7D*, despite their differences in the LRP. The *Eps-7D*-*early* had faster rate of development of each individual floret and similar duration of floral development (due to an early onset of floret development) and that determined for distal florets to improve their rate of survival producing fertile florets. On the other hand, the *Eps-7D*-*late* delayed the initiation of distal florets which then developed at a slower rate, consequently diminishing the likelihood of survival of many of the labile florets primordia, reaching anthesis with less fertile florets, overriding the effect of bearing slightly higher spikelet number per spike on the overall number of fertile florets per spike. The effect of *Eps-2B* on spike fertility was in alignment with the general hypothesis mentioned above in that the *Eps-2B*-*late* allele improved spike fertility (when the allele of *Eps-7D* in the background was *late*) due to a slightly longer duration of floret development resulting in lower floret mortality as evidenced with other *Eps* genes^36^; and through extending the LRP manipulating the photoperiod^13,41,42^. Our results (when spike fertility was higher) further provide proof to the concept that variations in the floret fertility are mainly due to differences in the survivability (low mortality) of produced florets (due to variations in duration, rate of development or resource availability) rather than improved maximum number of florets initiated^41,43^.

## Materials and methods

### Plant material, field conditions and management

Two more field experiments were carried out in two consecutive cropping seasons, 2016-17 (CS1) and 2017-18 (CS2) under similar environmental conditions near Bell-lloc d’Urgell (Lat. 41º38′ N, 0º44′ E in CS1 and Lat. 41º37′ N, 0º47′ E in CS2), Lleida, North-East Spain. Both experiments were sown within optimum sowing dates (16 November 2016 and 17 November 2017). The seeds were sown at a density of 125 kg ha^-1^ with the aim of attaining a uniform plant density of 250 plants per m^2^. The experiments were maintained under stress-free conditions (i.e. weeds, insects, and diseases were controlled or prevented, and plots were irrigated and fertilized as required).

Weather data were collected daily from Meteocat (agro-meteorological network of Catalonia, en.meteocat.gencat.cat) from a station located near the experimental site. The average temperatures from sowing to maturity (growing period) were 12.6 and 11.8 ºC (Fig. 11a) and accumulated precipitations for the whole growing period were 263.6 and 286.9 mm for CS1 and CS2, respectively (Fig. 11b). With respect to average temperature, the two growing seasons were slightly colder in the early phase (sowing to 2 leaf stage) but very similar during early and late reproductive phases until maturity when compared to the average of the last five years (2010-2016).

### Treatments and experimental design

Treatments consisted of eight lines (Table 4) derived from the cross Paragon × Baj where Paragon and Baj carry late and early allele of both *Eps-7D* and *Eps-2B*, respectively. Although not NILs, lines mostly differed for these two *Eps* QTLs (7D and 2B) while having similar genetic composition (mixture of Paragon and Baj) for the other major genes influencing anthesis time. The heading date QTL identified in Paragon x Baj are shown in Table 1 and QTL plots in supplementary Fig. S1. RILs chosen for detailed study were fixed for *Ppd-D1* and *Rht-B1*, which are both segregating in this population. Combining the different lines for their alleles of these two *Eps* genes, we end up with four contrasts to test the direct effects of these alleles as well as the interactions between them (i.e. to what degree the allelic form of one of these genes may condition, qualitatively or quantitatively, the effects of the other one).

**Table 4.** Lines selected for this study, possessing contrasting alleles of both *Eps* genes (*Eps-7D* and *Eps-2B*). For comparing the effects of these alleles we averaged the results across lines possessing the same alleles of these two genes (see scheme for contrasts on the right and footnote). That offered four contrasting combinations of the two alleles (*late or early*) of the two *Eps* genes; and depending on the particular comparisons made the direct and interactive effects of these genes were studied.

| Line | *Eps* allele (*source*) |  | Contrasts |
| --- | --- | --- | --- |
|  | *7D* | *2B* |  |
| Paragon x Baj 19 | *late* (Paragon) | *late* (Paragon) | *Eps-7D-late* + *Eps-2B-late* |
| Paragon x Baj 34 | *late* (Paragon) | *early* (Baj) | *Eps-7D-late* + *Eps-2B-early* |
| Paragon x Baj 93 | *late* (Paragon) | *early* (Baj) |  |
| Paragon x Baj 107 | *late* (Paragon) | *early* (Baj) |  |
| Paragon x Baj 132 | *late* (Paragon) | *early* (Baj) |  |
| Paragon x Baj 48 | *early* (Baj) | *late* (Paragon) | *Eps-7D-early* + *Eps-2B-late* |
| Paragon x Baj 104 | *early* (Baj) | *early* (Baj) | *Eps-7D-early* + *Eps-2B-early* |
| Paragon x Baj 147 | *early* (Baj) | *early* (Baj) |  |

1 and 2.- Direct effects of *Eps-7D* and *Eps-2B* genes, respectively.
3 and 4.- Effects of *Eps-7D* gene when the background has the *late* and *early* allele of *Eps-2B* gene, respectively.
5 and 6.- Effects of *Eps-2B* gene when the background has the *late* and *early* allele of *Eps-7D* gene, respectively.

Genotypes were arranged in a randomized complete block design with three blocks. Plot size was six rows (0.20 m apart) wide and 4 m long. Three plants in each plot (9 plants per treatment) were marked after the seedling emergence stage. These marked plants were visually representative of the whole plot as well as had uniform plant density around them and were randomly chosen from any of the 4 central rows. In addition, we marked 1 linear meter in each experimental unit which showed uniform emergence of seedlings and optimum plant-to-plant density to be sampled for measurements at anthesis.

## Measurements and data analysis

### Genetic mapping and QTL analysis

The Paragon x Baj RILs were genotyped using the 35k Axiom Wheat Breeders’ array (Affymetrix^44^) and a genetic map was constructed using MSTmap online (http://mstmap.org/) with default setting and a grouping threshold of LOD = 10. Linkage groups were separated into different chromosomes manually. Full genotype files and genetic maps can be found at https://data.cimmyt.org/dataset.xhtml?persistentId=hdl:11529/10996. Employing the genetic map, QTL detection was performed on heading date measurements from the UK trial, using package “qtl” (vs. 1.44–9-^45^) in two st*Eps*, the first scan determining co-factors and the second scan identifying robust QTL, taking the co-factors into account^46^.

### Leaf number and dynamics of leaf appearance

The number of leaves emerging from the main shoot was recorded in the three marked plants once or twice a week depending on temperature. Measurements began from 1 leaf stage and continued until the flag leaf was fully emerged, using the scale described by Haun (1973)^47^. Leaf appearance dynamics were analyzed by regressing the number of leaves emerged against the calculated thermal time (using daily average temperature and assuming a base temperature of 0 ºC) from sowing. Despite the fact that the linear regressions in all cases were highly significant, the distribution of residuals indicated that a bi-linear trend was required (to have a random distribution of residuals). Therefore, we fitted a segmented linear regression forcing a breakpoint in rate of leaf appearance at leaf 7, because it has been shown that (i) this change in phyllochron coincides with Haun stages between 6 and 8^12,48–50^ and (ii) using that threshold for the change in slope of the bi-linear regression produces excellent outputs in a wide range of environmental (e.g.^14^) and genotypic contexts (e.g.^51,52^). Phyllochron is the time between the appearance of two successive leaves. It was calculated as the reciprocal of the rate of leaf appearance. In each case two values are generated, corresponding to the early (leaves 1-7, phyllochron I) and late appearing leaves (leaf 7-flag leaf, phyllochron II).

### Shoot apex dissection and dynamics of primordia initiation

To determine the apical stage and number of primordia in the main shoot apex, one representative plant per experimental unit (three plants per treatment) was sampled randomly from the central rows. This was performed once or twice a week, depending on temperature, from the 2-leaf stage to terminal spikelet stage. The plants were taken to the laboratory for dissection under a binocular microscope (Leica MZ 8.0, Leica Microsystems, Heerbrugg, Switzerland). Apical stages were determined using the scale described by Kirby and Appleyard^53^. Time taken to reach apical stages including double ridge and terminal spikelet (TS) were recorded for all the lines in each block. Along with the apical stages, the number of leaf and spikelet primordia initiated at each sampling were counted. After TS stage, one plant from each experimental unit was sampled two or three times a week, again depending on temperature, until anthesis. This was to count the number of florets initiated and determine the stage of development for each floret primordium following the scale developed by Waddington et al. (1983)^54^. Wheat exhibits asynchronous development and growth of spikelets within the spike and florets within the spikelets is different. For that reason, we dissected 3 spikelets per spike, one from apical (3^rd^ or 4^th^ from the top) and one from central and one from the basal (3^rd^ or 4^th^ from the bottom) position of the spike. The florets within each spikelet position were numbered from F1 to Fn according to their position with respect to rachis, where F1 being the floret that was most proximal to the rachis and Fn the most distal floret. Floret development was observed from an early floret primordia stage (<W3.5) to until W10 stage (stage when floret is already fertile) or highest stage attained by florets that was aborted. The time taken for the F1 to reach W10 stage from sowing was considered as the anthesis time and late reproductive phase (LRP) was calculated as period between TS and anthesis time.

### Living floret primordia and fertile floret number

The number of living floret primordia was derived from the individual floret (F1 to Fn) development curve. The sum of all the florets that continued to develop was calculated for each sampling date, and then plotted against thermal time. This shows the maximum number of florets initiated and final number of fertile florets at anthesis. Only florets that reached W4.5 (stage when stamen, pistil and carpel primordia are present) were considered for calculating living floret primordia. While F1 and F2 florets invariably (in all genotypes and spikelet positions) reached W10, fate of F3 to F6 florets was depended on genotype, spikelet position and field conditions but these florets at least reached W4.5 in all cases. Florets F7-F10 were initiated in most cases, but did not reach the threshold of W4.5 stage in most spikelets.

### Fertile floret mapping at anthesis

Three plants were randomly selected from the marked 1 m of uniform plants in each experimental unit (9 plants per treatment) to “map” the number of fertile florets per spike, through counting them at each spikelet in the main-shoot spike (i.e. the number of fertile florets was mapped for every spikelet position from basal to terminal spikelet; S1 to Sn). Florets were considered fertile if they were at least at the green anther stage (>W8.5), considering the florets within and across spikelets have asynchronous development and that floret death is highly unlikely when florets have progressed in development to very close to the stage of fertile floret.

### Analysis

The data of the variables studied here was subjected to analysis of variance using the statistical software JMP® Pro Version 14.0 (SAS Institute Inc. Cary, NC, USA). The significance level for the differences between Eps allele were calculated using LSmMean contrast. The two-way interaction between two Eps QTLs and three-way interaction between Eps QTLs and season were analyzed using full factorial model in JMP® Pro version 14.0.

## Acknowledgements

Funding was partly provided by projects AGL2015-69595-R, from the Spanish Research Agency (AEI), and IWYP25FP, from the International Wheat Yield Partnership (IWYP). We thank Semillas Batlle for their technical support in maintenance of field experiments. We are indebted with Prof. Ignacio Romagosa for helping with the statistical analyses. PB held a pre-doctoral research contract from the agency for Management of University and Research (AGAUR) from the *Generalitat de Catalunya*.

## Notes

### Competing Interest Statement

The authors have declared no competing interest.

## References

1. Snape, J. W., Butterworth, K., Whitechurch, E. & Worland, A. J. Waiting for fine times: Genetics of flowering time in wheat. Euphytica 119, 185–190 (2001).

2. Kamran, A., Iqbal, M. & Spaner, D. Flowering time in wheat (Triticum aestivum L.): A key factor for global adaptability. Euphytica 197, 1–26 (2014).

3. Richards, R. A. Crop improvement for temperate Australia: Future opportunities. Field Crops Res. 26, 141–169 (1991).

4. Zheng, B., Chapman, S. C., Christopher, J. T., Frederiks, T. M. & Chenu, K. Frost trends and their estimated impact on yield in the Australian wheatbelt. J. Exp. Bot. 66, 3611–3623 (2015).

5. Araus, J. L., Slafer, G. A., Reynolds, M. P. & Royo, C. Plant breeding and drought in C3 cereals: What should we breed for? Ann. Bot. 89, 925–940 (2002).

6. Worland, A. J. The influence of flowering time genes on environmental adaptability in European wheats. Euphytica 89, 49–57 (1996).

7. Nazim Ud Dowla, M.A.N., Edwards, I., O’Hara, G., Islam, S. & Ma, W. Developing Wheat for Improved Yield and Adaptation Under a Changing Climate: Optimization of a Few Key Genes. Engineering 4, 514–522 (2018).

8. Craufurd, P. Q. & Wheeler, T. R. Climate change and the flowering time of annual crops. J. Exp. Bot. 60, 2529–2539 (2009).

9. Semenov, M. A., Stratonovitch, P., Alghabari, F. & Gooding, M. J. Adapting wheat in Europe for climate change. J. Cereal Sci. 59, 245–256 (2014).

10. Slafer, G. A. & Rawson, H. M. Sensitivity of wheat phasic development to major environmental factors: A re-examination of some assumptions made by physiologists and modellers. Aust. J. Plant Physiol. 21, 393–426 (1994).

11. Whitechurch, E. M. & Slafer, G. A. Contrasting Ppd alleles in wheat: Effects on sensitivity to photoperiod in different phases. Field Crops Res. 73, 95–105 (2002).

12. González, F. G., Slafer, G. A. & Miralles, D. J. Pre-anthesis development and number of fertile florets in wheat as affected by photoperiod sensitivity genes Ppd-D1 and Ppd-B1. Euphytica 146, 253–269 (2005).

13. Miralles, D. J., Richards, R. A. & Slafer, G. A. Duration of the stem elongation period influences the number of fertile florets in wheat and barley. Funct. Plant Biol. 27, 931 (2000).

14. Slafer, G. A., Abeledo, L. G., Miralles, D. J., Gonzalez, F. G. & Whitechurch, E. M. Photoperiod sensitivity during stem elongation as an avenue to raise potential yield in wheat. Euphytica 119, 191–197 (2001).

15. Rawson, H. M., Zajac, M. & Penrose, L. D. J. Effect of seedling temperature and its duration on development of wheat cultivars differing in vernalization response. Field Crops Res. 57, 289–300 (1998).

16. Slafer, G. A. & Rawson, H. M. Responses to photoperiod change with phenophase and temperature during wheat development. Field Crops Res. 46, 1–13 (1996).

17. Whitechurch, E. M. & Snape, J. W. Developmental responses to vernalization in wheat deletion lines for chromosomes 5A and 5D. Plant Breed. 122, 35–39 (2003).

18. Allard, V. et al. The quantitative response of wheat vernalization to environmental variables indicates that vernalization is not a response to cold temperature. J. Exp. Bot. 63, 847–857 (2012).

19. Shaw, L. M., Turner, A. S. & Laurie, D. A. The impact of photoperiod insensitive Ppd-1a mutations on the photoperiod pathway across the three genomes of hexaploid wheat (Triticum aestivum). Plant J. 71, 71–81 (2012).

20. Pérez-Gianmarco, T. I., Slafer, G. A. & González, F. G. Wheat pre-Anthesis development as affected by photoperiod sensitivity genes (Ppd-1) under contrasting photoperiods. Funct. Plant Biol. 45, 645–657 (2018).

21. Griffiths, S. et al. Meta-QTL analysis of the genetic control of ear emergence in elite European winter wheat germplasm. Theor. Appl. Genet. 119, 383–395 (2009).

22. Steinfort, U., Trevaskis, B., Fukai, S., Bell, K. L. & Dreccer, M. F. Vernalisation and photoperiod sensitivity in wheat: Impact on canopy development and yield components. Field Crops Res. 201, 108–121 (2017).

23. Kamran, A. et al. Earliness per se QTLs and their interaction with the photoperiod insensitive allele Ppd-D1a in the Cutler × AC Barrie spring wheat population. Theor. Appl. Genet. 126, 1965–1976 (2013).

24. Gomez, D. et al. Effect of Vrn-1, Ppd-1 genes and earliness per se on heading time in Argentinean bread wheat cultivars. Field Crops Res. 158, 73–81 (2014).

25. Alvarez, M. A., Tranquilli, G., Lewis, S., Kippes, N. & Dubcovsky, J. Genetic and physical mapping of the earliness per se locus Eps-A m 1 in Triticum monococcum identifies EARLY FLOWERING 3 (ELF3) as a candidate gene. Funct. Integr. Genomics 16, 365–382 (2016).

26. Zikhali, M. et al. Validation of a 1DL earliness per se (eps) flowering QTL in bread wheat (Triticum aestivum). Mol. Breed. 34, 1023–1033 (2014).

27. Lewis, S., Faricelli, M. E., Appendino, M. L., Valárik, M. & Dubcovsky, J. The chromosome region including the earliness per se locus Eps-A m1 affects the duration of early developmental phases and spikelet number in diploid wheat. J. Exp. Bot. 59, 3595–3607 (2008).

28. Sukumaran, S. et al. Identification of earliness per se flowering time locus in spring wheat through a genome-wide association study. Crop Sci. 56, 2962–2972 (2016).

29. Zikhali, M. & Griffiths, S. The Effect of Earliness per se (Eps) Genes on Flowering Time in Bread Wheat. in Advances in Wheat Genetics: From Genome to Field 339–345 (Springer Japan, 2015). doi:10.1007/978-4-431-55675-6_39.

30. Slafer, G. A. Genetic basis of yield as viewed from a crop physiologist’s perspective. Ann. Appl. Biol. 142, 117–128 (2003).

31. Slafer, G. A. & Rawson, H. M. Intrinsic earliness and basic development rate assessed for their response to temperature in wheat. Euphytica 83, 175–183 (1995).

32. Le Gouis, J. et al. Genome-wide association analysis to identify chromosomal regions determining components of earliness in wheat. Theor. Appl. Genet. 124, 597–611 (2012).

33. Ochagavía, H., Prieto, P., Zikhali, M., Griffiths, S. & Slafer, G. A. Earliness Per Se by Temperature Interaction on Wheat Development. Sci. Rep. 9, 2584 (2019).

34. Ejaz, M. & von Korff, M. The genetic control of reproductive development under high ambient temperature. Plant Physiol. 173, 294–306 (2017).

35. Ochagavía, H., Prieto, P., Savin, R., Griffiths, S. & Slafer, G. A. Earliness per se effects on developmental traits in hexaploid wheat grown under field conditions. Eur. J. Agron. 99, 214–223 (2018).

36. Prieto, P., Ochagavía, H., Savin, R., Griffiths, S. & Slafer, G. A. Physiological determinants of fertile floret survival in wheat as affected by earliness per se genes under field conditions. Eur. J. Agron. 99, 206–213 (2018).

37. Bullrich, L., Appendino, M. L., Tranquilli, G., Lewis, S. & Dubcovsky, J. Mapping of a thermo-sensitive earliness per se gene on Triticum monococcum chromosome 1Am. Theor. Appl. Genet. 105, 585–593 (2002).

38. Appendino, M. L. & Slafer, G. A. Earliness per se and its dependence upon temperature in diploid wheat lines differing in the major gene Eps-A m 1 alleles. J. Agric. Sci. 149–154 (2003)

39. Miralles, D. J. & Slafer, G. A. Sink limitations to yield in wheat: how could it be reduced? J. Agric. Sci. 145, 139 (2007).

40. Hoogendoorn, J. The physiology of variation in the time of ear emergence among wheat varieties from different regions of the world. Euphytica 34, 559–571 (1985).

41. Serrago, R. A., Miralles, D. J. & Slafer, G. A. Floret fertility in wheat as affected by photoperiod during stem elongation and removal of spikelets at booting. Eur. J. Agron. 28, 301–308 (2008).

42. Prieto, P., Ochagavía, H., Savin, R., Griffiths, S. & Slafer, G. A. Dynamics of floret initiation/death determining spike fertility in wheat as affected by Ppd genes under field conditions. J. Exp. Bot. 69, 2633–2645 (2018).

43. Miralles, D. J., Katz, S. D., Colloca, A. & Slafer, G. A. Floret development in near isogenic wheat lines differing in plant height. Field Crops Res. 59, 21–30 (1998).

44. Allen, A. M. et al. Characterization of a Wheat Breeders’ Array suitable for high-throughput SNP genotyping of global accessions of hexaploid bread wheat (Triticum aestivum). Plant Biotechnol. J. 15, 390–401 (2017).

45. Broman, K. W., Wu, H., Sen, Ś. & Churchill, G. A. R/qtl: QTL mapping in experimental crosses. Bioinformatics 19, 889–890 (2003).

46. Wingen, L. U. et al. Wheat landrace genome diversity. Genetics 205, 1657–1676 (2017).

47. Haun, J. R. Visual Quantification of wheat development. Agron. J. 65, 116–119 (1973).

48. Jamieson, P. D., Brooking, I. R., Porter, J. R. & Wilson, D. R. Prediction of leaf appearance in wheat: a question of temperature. Field Crops Res. 41, 35–44 (1995).

49. Slafer, G. A. & Rawson, H. M. Phyllochron in wheat as affected by photoperiod under two temperature regimes. Aust. J. Plant Physiol. 24, 151–158 (1997).

50. Calderini, D. F., Miralles, D. J. & Sadras, V. O. Appearance and growth of individual leaves as affected by semidwarfism in isogenic lines of wheat. Ann. Bot. 77, 583–589 (1996).

51. Ochagavía, H., Prieto, P., Savin, R., Griffiths, S. & Slafer, G. A. Duration of developmental phases, and dynamics of leaf appearance and tillering, as affected by source and doses of photoperiod insensitivity alleles in wheat under field conditions. Field Crops Res. 214, 45–55 (2017).

52. Ochagavía, H., Prieto, P., Savin, R., Griffiths, S. & Slafer, G. A. Dynamics of leaf and spikelet primordia initiation in wheat as affected by Ppd-1a alleles under field conditions. J. Exp. Bot. 69, 2621–2631 (2018).

53. Kirby, E. J. M. & Appleyard, M. Development and structure of the wheat plant. in Wheat Breeding 287–311 (Springer Netherlands, 1987).

54. Waddington, S. R., Cartwright, P. M. & Wall, P. C. A Quantitative Scale of Spike Initial and Pistil Development in Barley and Wheat. Ann. Bot. 51, 119–130 (1983).

