## Supplementary for "Interactions between two QTLs for time to anthesis on spike development and fertility in wheat"

**Supplementary data**

### Supplementary data

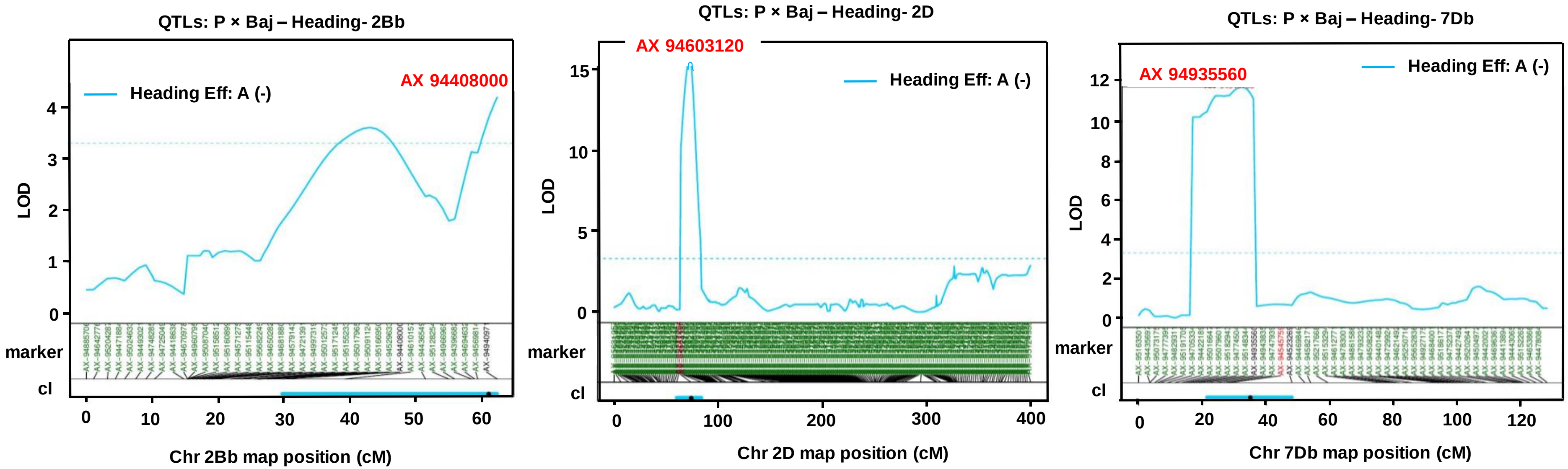


**Supplementary Figure S1.** Heading date QTL on chromosome 2B, 2D and 7D. LOD scores are plotted along the chromosome axis. Marker names and position are listed underneath, along the chromosome axis. The peak marker is highlighted in red and the markers bordering the confidence interval in black. The extend of the confidence interval is shown as horizontal blue lines underneath the marker names.

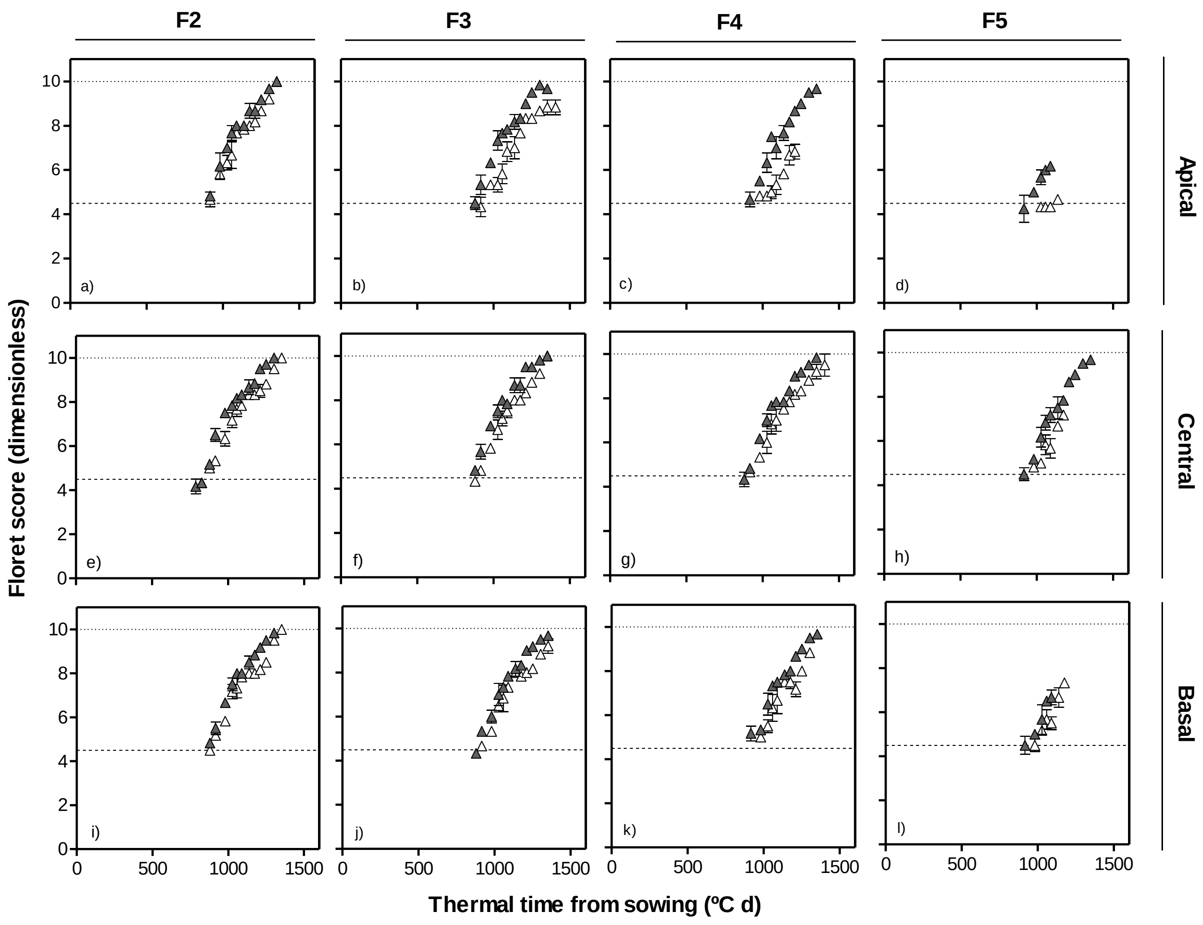


**Supplementary Figure S2.** Dynamics of floret development (dimensionless floret score) in F2, F3, F4 and F5 florets at apical (top panel: a-d), central (middle panel: e-h) and basal (bottom panel: i-l) positions of spike with thermal time from sowing in lines with *Eps-7D-late* (open symbol) and early (closed symbol) allele with the late allele of *Eps-2B* in the background in second cropping season.

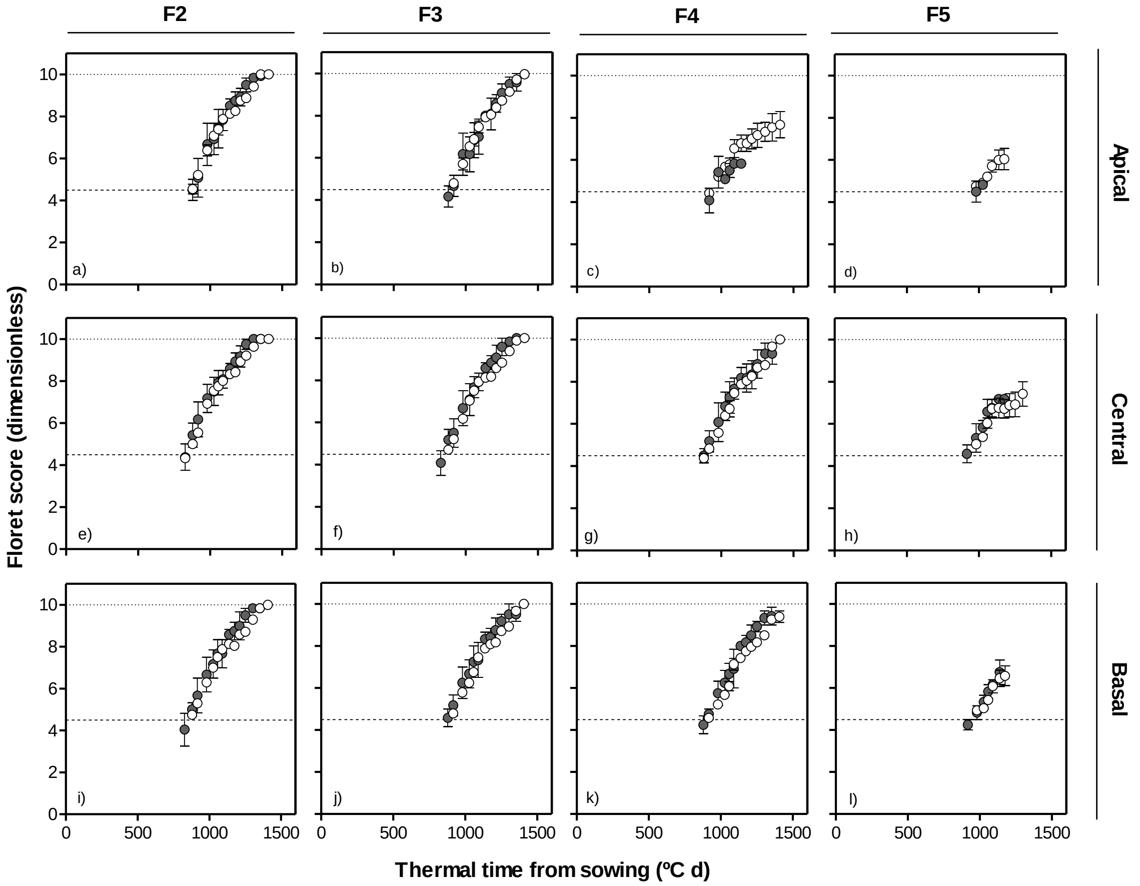


**Supplementary Figure S3.** Dynamics of floret development (dimensionless floret score) in F2, F3, F4 and F5 florets at apical (top panel: a-d), central (middle panel: e-h) and basal (bottom panel: i-l) positions of spike with thermal time from sowing in lines with *Eps-7D-late* (open symbol) and early (closed symbol) allele with the early allele of *Eps-2B* in the background in second cropping season.

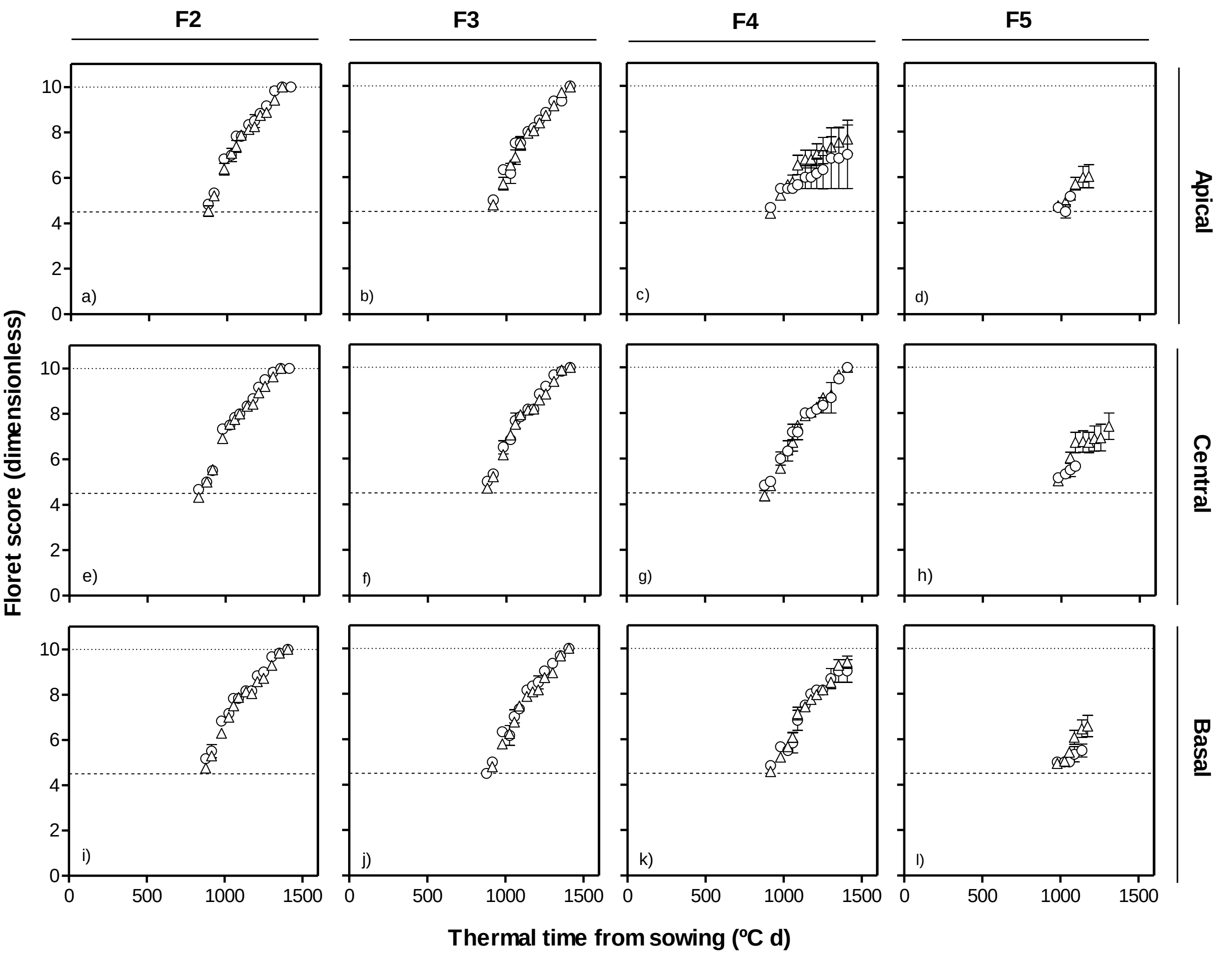


**Supplementary Figure S4.** Dynamics of floret development (dimensionless floret score) in F2, F3, F4 and F5 florets at apical (top panel: a-d), central (middle panel: e-h) and basal (bottom panel: i-l) positions of spike with thermal time from sowing in lines with *Eps-2B-late* (triangles) and *early* (circles) allele with the *late* allele of *Eps-7D* in the background in second cropping season.

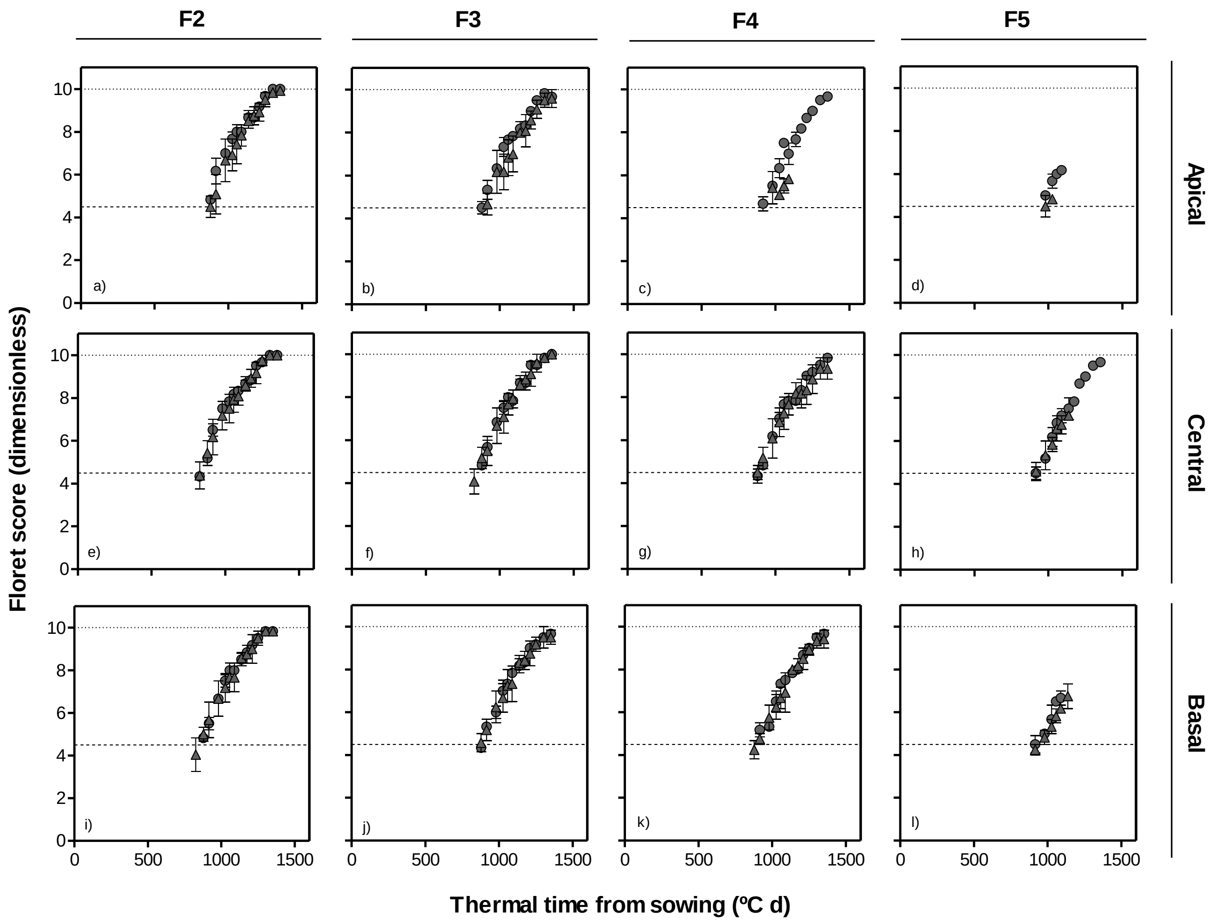


**Supplementary Figure S5.** Dynamics of floret development (dimensionless floret score) in F2, F3, F4 and F5 florets at apical (top panel: a-d), central (middle panel: e-h) and basal (bottom panel: i-l) positions of spike with thermal time from sowing in lines with *Eps-2B-late* (triangles) and *early* (circles) allele with the *early* allele of *Eps-7D* in the background in second cropping season.

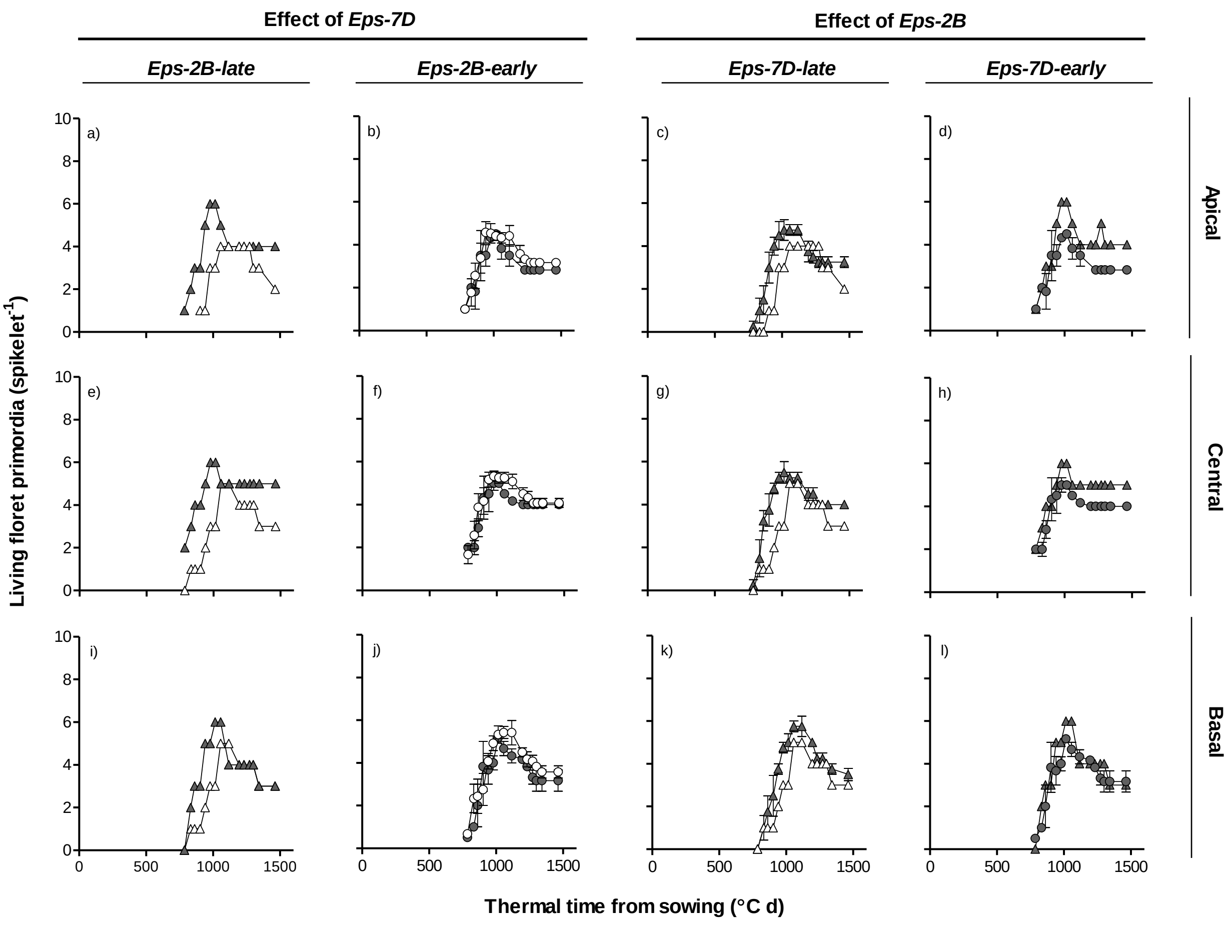


**Supplementary Figure S6.** Number living floret primordia at apical (top panels: a-d) central (middle panel: e-h) and basal spikelet (bottom panel: i-l) and thermal time from sowing as affected by *Eps-7D* (left panels: a, b, e, f, i and j) and *Eps-2B* genes (right panels: c, d, g, h, k and i) on backgrounds contrasting in the allelic form of the other *Eps* gene (left and right panels within each *Eps* gene) in first cropping season.

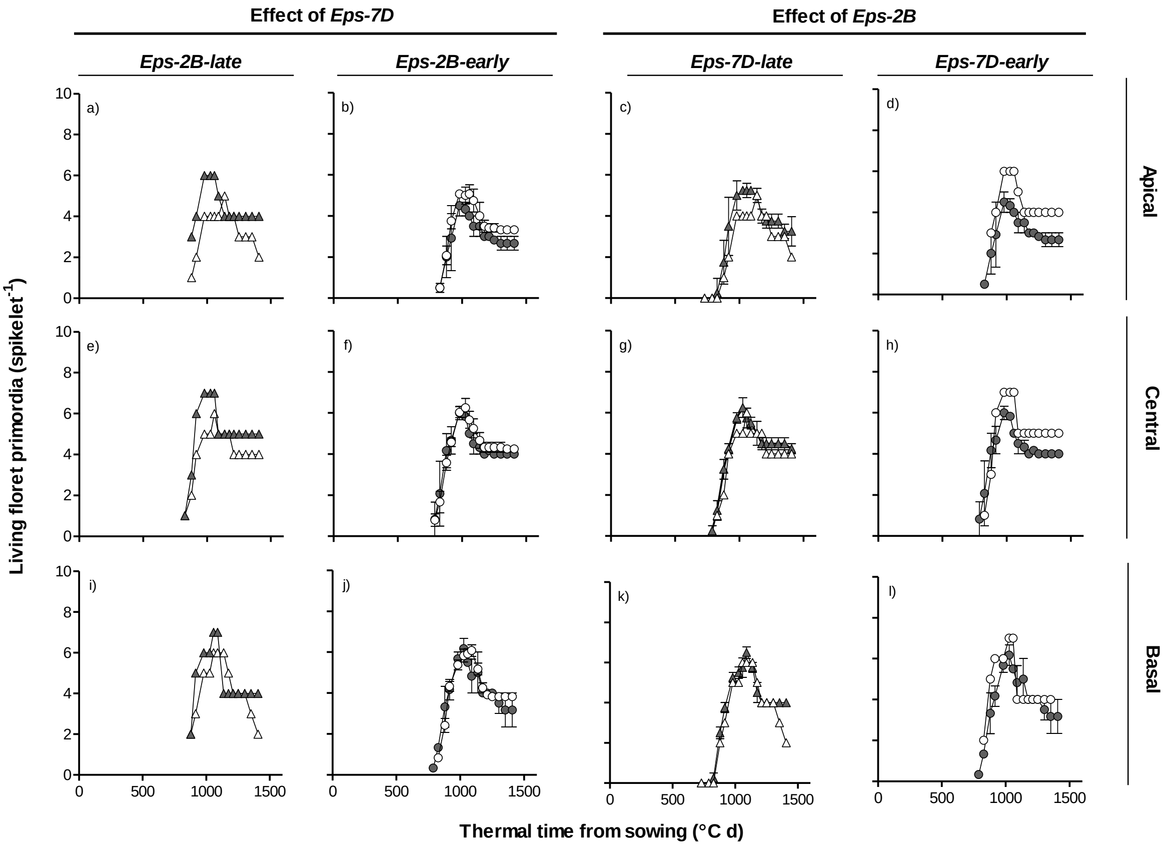


**Supplementary Figure S7.** Number living floret primordia at apical (top panels: a-d) central (middle panel: e-h) and basal spikelet (bottom panel: i-l) and thermal time from sowing as affected by *Eps-7D* (left panels: a, b, e, f, i and j) and *Eps-2B* genes (right panels: c, d, g, h, k and i) on backgrounds contrasting in the allelic form of the other *Eps* gene (left and right panels within each *Eps* gene) in second cropping season.
